# MR-AIV reveals *in vivo* brain-wide fluid flow with physics-informed AI

**DOI:** 10.1101/2025.07.30.667741

**Authors:** Juan Diego Toscano, Yisen Guo, Zhibo Wang, Mohammad Vaezi, Yuki Mori, George Em Karniadakis, Kimberly A.S. Boster, Douglas H. Kelley

## Abstract

The circulation of cerebrospinal and interstitial fluid plays a vital role in clearing metabolic waste from the brain, and its disruption has been linked to neurological disorders. However, directly measuring brain-wide fluid transport—especially in the deep brain—has remained elusive. Here, we introduce magnetic resonance artificial intelligence velocimetry (MR-AIV), a framework featuring a specialized physics-informed architecture and optimization method that reconstructs three-dimensional fluid velocity fields from dynamic contrast-enhanced magnetic resonance imaging (DCE-MRI). MR-AIV unveils brain-wide velocity maps while providing estimates of tissue permeability and pressure fields—quantities inaccessible to other methods. Applied to the brain, MR-AIV reveals a functional landscape of interstitial and perivascular flow, quantitatively distinguishing slow diffusion-driven transport (∼ 0.1 µm/s) from rapid advective flow (∼ 3 µm/s). This approach enables new investigations into brain clearance mechanisms and fluid dynamics in health and disease, with broad potential applications to other porous media systems, from geophysics to tissue mechanics.

## 1. Introduction

Fluids permeate the brain. Their circulation clears metabolic wastes whose accumulation is linked to diseases like Alzheimer’s and is altered in pathological conditions like stroke [1–5]. Mapping and quantifying the flow of cerebrospinal fluid (CSF) in large spaces and the putative flow of interstitial fluid (ISF) in small extracellular spaces (ECSs) is crucial to understanding the function, failure, and potential rehabilitation of the brain’s waste removal, or “glymphatic,” system. Yet, direct measurement of the brain-wide fluid movement has been impossible. This inability to quantify the fluid dynamics governing waste clearance has fundamentally limited our understanding of diseases like Alzheimer’s.

Current techniques for imaging brain flows are often invasive, superficial, or lack spatial resolution. Tracking particles imaged in vivo with two-photon microscopy [4, 6–8] yields high-fidelity velocity measurements (especially when supplemented with artificial intelligence velocimetry [9, 10]), but can be done only in regions near the brain surface, such as pial (surface) perivascular spaces (PVSs) of mice. Flow velocities have been estimated by tracking tracer dye fronts in brain-wide imaging in mice [4, 11, 12], but front tracking neglects diffusion and out-of-plane flow. Both particle tracking and front tracking require invasive surgeries, like cranial windows or scalp removal, ruling out widespread clinical use with human patients. Dynamic contrast-enhanced magnetic resonance imaging (DCE-MRI) requires no surgery and measures signal from an injected tracer, over space and time. DCE-MRI has been used to quantify transport with apparent diffusion coefficients [13–15], but transport by bulk flow (advection) is a fundamentally different process than diffusion and cannot accurately be described by such coefficients. Data-driven approaches show promise for biological flow modelling [16–19], but without direct measurements of velocity in the deep brain, data-driven methods cannot be applied.

DCE-MRI has been combined with other methods to estimate velocities from observed tracer concentrations. Flows in rat brains have been mapped using optimal mass transport (OMT) [20–24], an optimal control approach that accounts for diffusion and noise using a regularization term. However, OMT uses a diffusivity-like constant in the mass transport equation that is heuristically chosen, not stated in terms of physical diffusion coefficients, making it difficult to compare the results with other works. Flows in human brains have been estimated by fitting several different models to DCE-MRI measurements using adjoint methods [25]. However, neither OMT nor adjoint methods have been validated with synthetic data. Additionally, neither can infer permeability or pressure because those quantities are absent from the governing equations employed. Though DCE-MRI can accurately image an injected tracer as it spreads through the brain, mapping and quantifying the flow that is complicit in that transport is difficult. Calculating the velocity field from tracer motion requires solving an ill-posed inverse problem in which boundary conditions are usually unknown. In vivo measurements are sparse: particle tracking produces measurements only at particles’ locations; front tracking produces measurements only at fronts’ locations; and MRI has spatial resolution coarser than the size of key structures like PVSs, causing partial-volume effects. Measurement noise undermines many methods for solving inverse problems. Efflux routes are not well-known [26]. Velocity magnitudes in the brain span several orders of magnitude, from ∼ 10^1^ µm/s measured in PVSs [6] to perhaps ∼ 10^−2^ µm/s or less expected in ECSs [27]. Tissue properties like permeability, which strongly affect flows, are also likely to span wide ranges. Finally, the physics that governs brain flow is not well-known. Some open spaces harbour viscous flow governed by the Stokes equation [28]. Most of the brain, however, is usually modelled as a porous medium, with momentum governed by Darcy’s law [29] or the Darcy-Brinkman equation [30]. Some studies have modelled tissue as poroelastic [31, 32]. Physics involving fractional derivatives may also be applicable [33].

To address these challenges, we developed Magnetic Resonance Artificial Intelligence Velocimetry (MR-AIV), a physics-informed machine learning (PIML) framework that infers three-dimensional velocity, pressure, and permeability fields solely from tracer concentration data. MR-AIV fundamentally advances the capabilities of standard PIML [34] to tackled ill-posed problems from noisy experimental multiscale data through three major innovations.

First, unlike standard single-network models [9, 10, 35–37], MR-AIV uses a modular architecture with four specialized networks to independently model pressure, permeability, the clean concentration signal, and its associated noise. This structure enables Darcy’s Law and the steady-state assumptions to be incorporated exactly, thereby reducing the number of constraints and simplifying training. Additionally, encoding Darcy’s law reduces the number of variables to model partially mitigating the problem’s ill-posedness. Notably, having a dedicated network for permeability is crucial for tackling the multi-scale nature of the velocities. Since permeability in the brain spans several orders of magnitude, this modular component acts as an adaptive map, enabling the model to accurately capture the sharp transitions between slow interstitial and rapid perivascular flow.

Second, it directly confronts noisy experimental data by modeling the measurement error as Gaussian noise. This assumption enables the use of a Negative Log-Likelihood objective, allowing one of the network modules to explicitly learn the space- and time-dependent noise, while another module learns the clean, denoised concentration signal. The governing physical laws are then constrained only on this clean signal, not the raw data, preventing the model from fitting to noise and significantly improving performance and robustness.

Finally, we introduce time-dependent residual-based attention (TD-RBA), a novel optimization method that addresses the challenge of physical residuals that vary by orders of magnitude over time. This method guides the optimizer using physics-based attention, effectively acting as a point-wise adaptive learning rate that is informed by the governing equations, ensuring that all phases of the tracer’s transport contribute meaningfully to the final solution.

These three innovations were developed specifically to address the challenges inherent in inferring velocity from measurements of tracer injected in a brain, but they represent important improvements to PIML that could be applied to any flow through porous media governed by Darcy’s Law, including flows in geophysics, tissue mechanics, thrombosis, and biomechanics.

To test MR-AIV, we simulated the spread of tracer injected into the cisterna magna of a mouse, a method common in experiments, using a realistic geometry of a mouse brain with permeabilities spanning four orders of magnitude, a challenging feature we expect in real brains. An MR-AIV model, trained only on the simulated concentration, accurately reproduced not only the concentration but also the underlying velocity field, which was not used for training.

Model accuracy depended little on the initial guess for the permeability of brain tissue, though permeability varies by orders of magnitude across anatomical regions. Quantifying model uncertainty, we found that high uncertainty, slow flow, and high error were strongly correlated. Next, we applied MR-AIV to DCE-MRI measurements of in vivo cisterna magna injections in five wild-type mice. We inferred three-dimensional velocities throughout the brain, including deep regions inaccessible to optical imaging.

MR-AIV offers a data-driven framework for modeling brain fluid transport without requiring direct velocity measurements, taking a step toward comprehensive, non-invasive characterization of glymphatic flow and its potential alterations in aging and neurological diseases. Although in this work we demonstrate MR-AIV on mouse brains, it could potentially be applied to human brains because DCE-MRI, which is already used in clinical settings, is non-invasive. Our results demonstrate that this method can reveal physiologically plausible transport patterns in vivo, providing new tools for studying cerebrospinal fluid dynamics and potentially informing diagnosis and treatment of neurodegenerative disorders. Because the MR-AIV framework is generalizable to other imaging modalities, organ systems, and even non-biological flows, it has wide-reaching implications beyond neuroscience.

## 2. Results

### Results

#### 2.1. MR-AIV reconstructs velocity fields from concentration observations

We injected gadobutrol into the cisterna magna of five wild-type mice and acquired DCE-MRI data. This yielded a time-dependent change in tracer intensity, or signal enhancement ratio (SER), which we normalized by its maximum value and multiplied by 100, using the result as a proxy for concentration *c* across the brain in three dimensions (Fig. 1(A), Fig. C.13).

**Figure 1.**
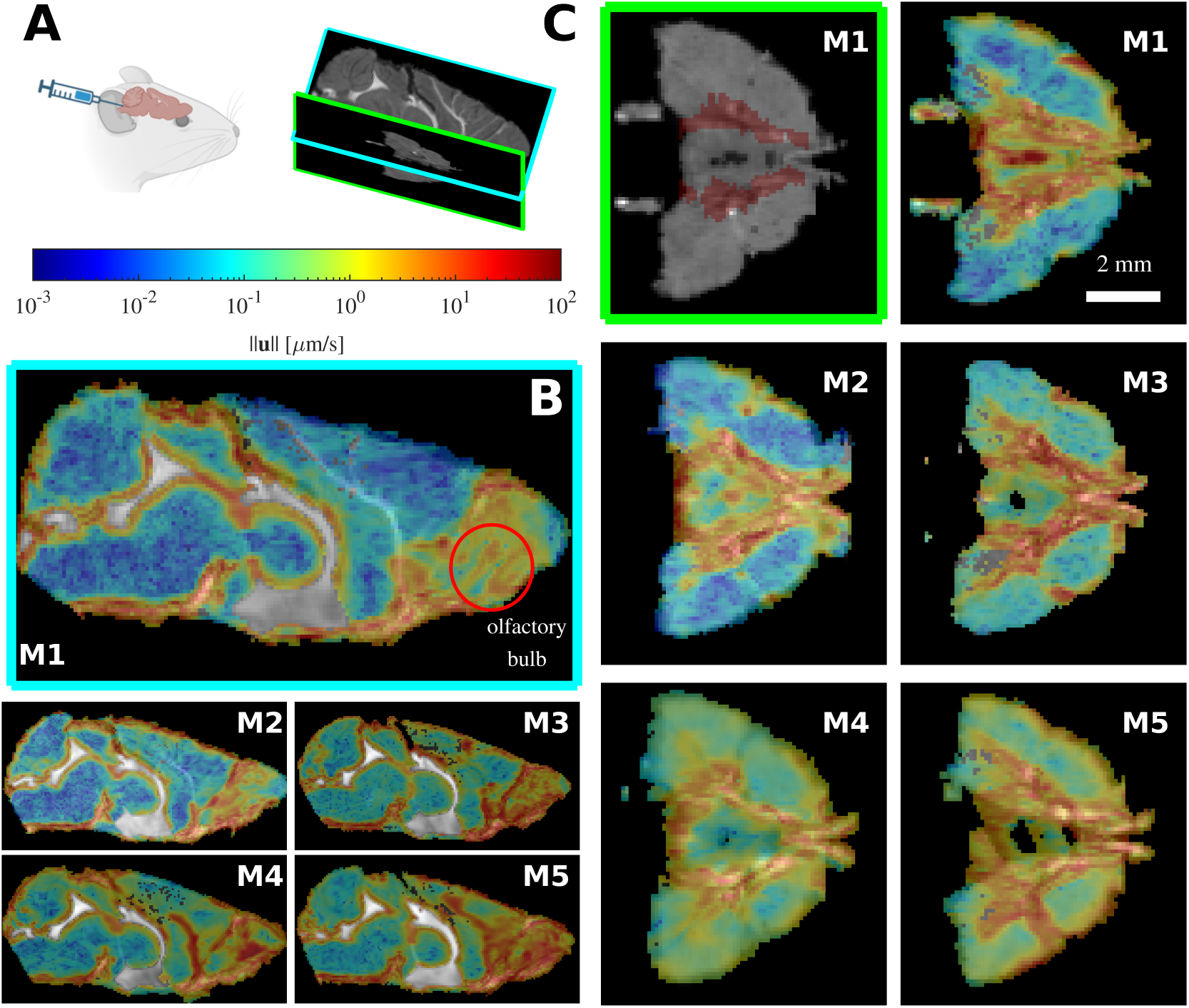
The MR-AIV inferred velocity magnitude || u || is similar across mice. (A) Gadobutrol is infused into the cisterna magna of five mice (M1–M5), and the tracer movement is recorded via dynamic contrast-enhanced magnetic resonance imaging. (B) The MR-AIV-inferred velocity magnitude in two planes (mid-sagittal, left, transverse plane, right). The Circle of Willis (location marked in red on the M1 structural image at upper left) can be seen in the transverse plane. Flow is consistently fast near the Circle of Willis and the olfactory bulb, which can be observed in the mid-sagittal plane. Velocity fields are overlaid on greyscale structural MRI images, which show through in excluded regions. The velocity magnitudes are similar for the five mice.

For each experiment, an MR-AIV model predicts the velocity **u** in the deep brain (Fig. 1). We observe a consistent structure across all five mice, as expected if anatomical brain structures shape the flow. Velocities are high in the olfactory bulb, near a well-known large subarachnoid space [38]. Chen et al. [23] also found high velocities in the olfactory bulbs of rats, using OMT. We also observe high velocities near the Circle of Willis, and in some mice, around the ventricles, suggesting flow between tissue and ventricles.

#### 2.2. MR-AIV reconstructs concentration fields from sparse observations

No previous measurements of flows deep in the brains of mice are available for comparison, so other assessments are required. One way to test MR-AIV is to compare the concentrations it reconstructs to the measured concentrations. We train our models on 50% of the concentration data, used both to learn the concentration field and to extract aleatoric uncertainty via the denoising module. Model performance is evaluated with the unseen data. In mice 1–5, the relative *L*^2^ error (see equation 3) was 8.63%, 9.07%, 12.21%, 13.14%, and 9.83%, respectively. Figure 2(A) shows the reconstructed concentration, point-wise error, and learned noise for Mouse 1 at three representative times. Predictions closely match measurements in magnitude and spatial variation. Notably, the absolute error appears unstructured, and the inferred uncertainty, trained to represent aleatoric noise, is spatially aligned with the absolute error. Most discrepancies seem to arise from measurement noise rather than modelling errors.

**Figure 2.**
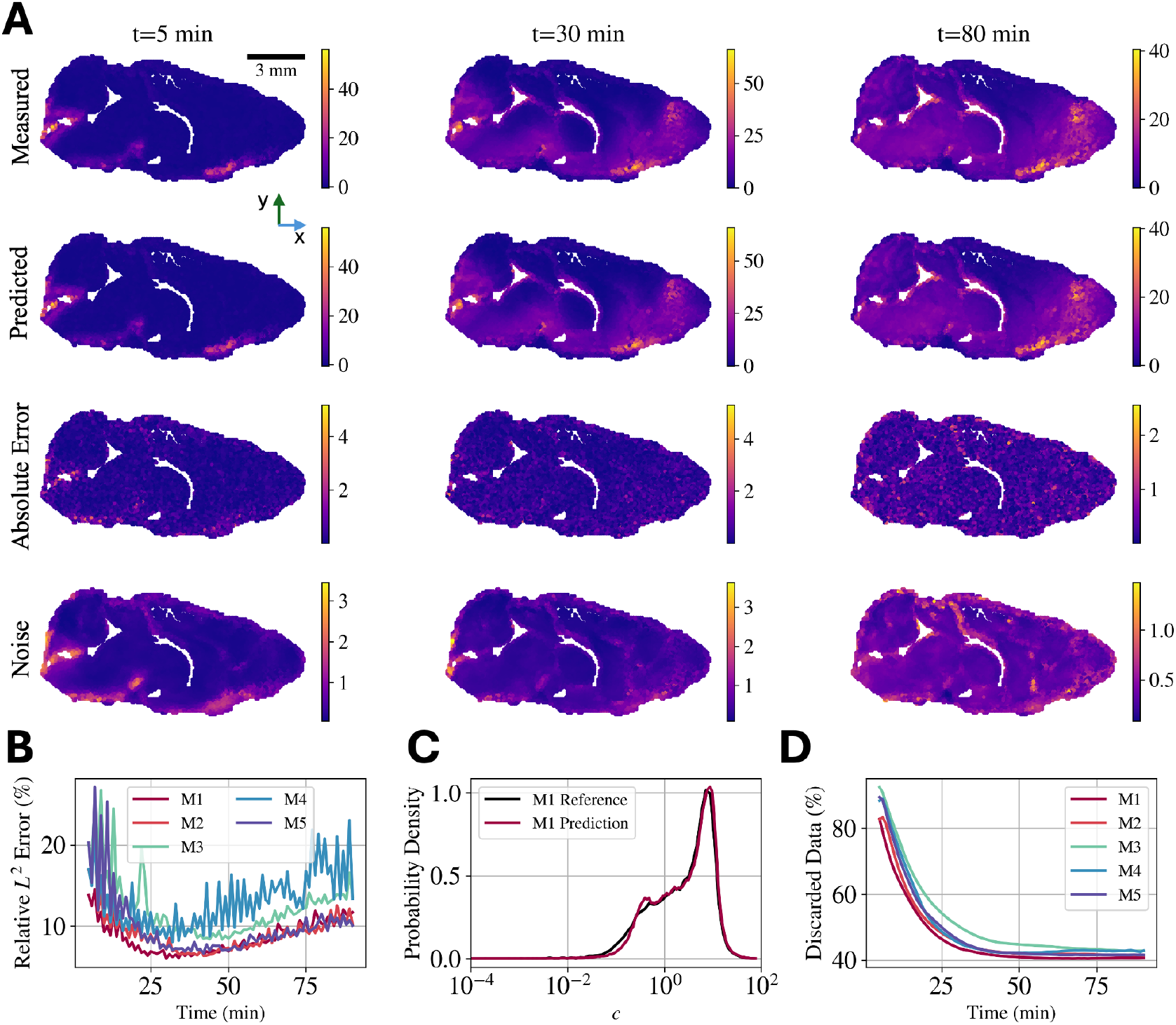
Concentration reconstruction using in-vivo data. (A) Measured concentration, reconstructed concentration, absolute error, and inferred noise for Mouse 1 (M1) on the mid-sagittal plane at *t* = 5, 30, and 80 minutes. The unstructured absolute error aligns spatially with the inferred noise, suggesting discrepancies are dominated by measurement uncertainty. (B) Relative *L*^2^ error over time for all five mice (M1–M5). The error drops sharply after the initial 10–20 minutes, averaging around 10%. (C) Probability density functions (PDFs, log scale) of measured and reconstructed concentration for M1. The model accurately captures the bimodal distribution, which spans six orders of magnitude. (D) Percentage of data discarded over time due to sensitivity thresholds. Data is discarded only from the PDE constraint during velocity inference, not from concentration training. Discard rates drop from 90% to ∼ 40% as the tracer spreads; higher discard rates correlate with larger reconstruction errors.

Figure 2(B) shows the temporal evolution of the relative *L*^2^ error for all five mice, which drops sharply after about 20 minutes, the time required for the tracer to reach a substantial fraction of the brain, and is steadier thereafter. Probability density functions (PDFs) of the reconstructed and experimental concentrations for Mouse 1 (Figure 2(C)) agree closely across six orders of magnitude. The concentration is bimodal.Most data from the first 20 minutes of an experiment are discarded because the concentration and its gradients are too small (Figure 2(D)). Once the tracer has spread, more measurements are retained, but about 40% of the brain is never reached by enough tracer to be useful for the MR-AIV model. Some brain regions are apparently perfused much less than others. Experiments where more data were discarded generally show larger relative *L*^2^ error.

#### 2.3. MR-AIV reveals bimodal velocity distributions across distinct brain regions

By inferring 3D velocity throughout the brain, MR-AIV makes detailed velocity statistics available. PDFs of velocity magnitude in all five mice are bimodal, with a low peak near 3 µm/s and a high peak on the order of 0.1 µm/s (Figure 3, top row). Bimodal distributions are consistent with the inferred velocity fields (Figure 1), which show slow flows in most of the brain but fast flows in a few regions. Individual velocity components showed similar distributions, though the peak near 3 µm/s was less distinct. By segmenting the inferred velocity maps according to the Allen Brain Atlas, a process whose accuracy stands to benefit from the continued development of next-generation atlases [39], we calculated velocity and permeability statistics for specific anatomical regions (Figure 3, Tables C.3, C.4, C.5, and C.6). Anatomical velocity variations were similar for all five mice, with slow flow in the hippocampus, caudate, thalamus, and sagittal sinus, but fast flow in the subarachnoid space near the olfactory bulb and near perivascular spaces adjacent to the Circle of Willis, middle cerebral artery, anterior cerebral artery, and basilar artery. Permeability tended to be high in the regions where flow was fast, and low in the regions where flow was slow (Fig. C.13(E)). According to Darcy’s law, velocity is the product of permeability and pressure gradient, so the observed similarity between velocity and permeability suggests that pressure gradients have similar magnitudes throughout the brain.

**Figure 3.**
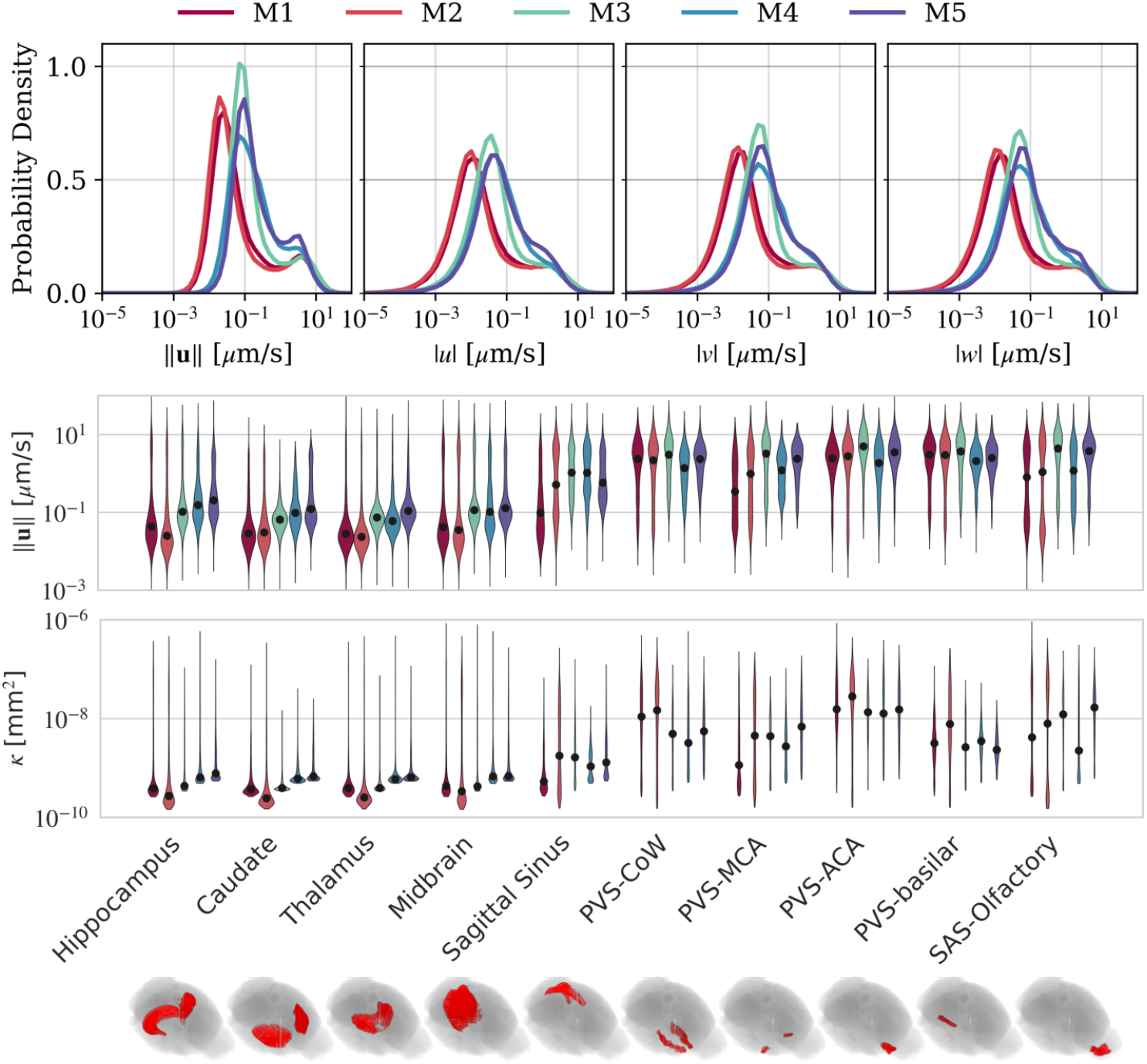
MR-AIV infers the three-dimensional velocity field throughout the brain. Probability density functions of velocity components (*u, v, w*) and velocity magnitude ||**u**|| are similar for five mice (M1-M5). The inferred velocity magnitude and permeability distributions in select anatomical regions. The median value is indicated with a black dot. Flows in the deep brain (e.g., hippocampus, caudate, thalamus, and midbrain) are slower. Faster flow and higher inferred permeability values are observed in the perivascular spaces (PVS) of the sagittal sinus, Circle of Willis (COW), middle cerebral artery (MCA), anterior cerebral artery (ACA), basilar artery, and subarachnoid space of the olfactory bulb.

#### 2.4. MR-AIV delineates advection- and diffusion-dominated transport

An important question for brain clearance concerns the relative importance of advection and diffusion in transporting solutes. Since MR-AIV infers both concentration and velocity fields, the advective and diffusive transport terms can be calculated at any point in space and time (see equation 1). Their time-averaged ratio is a local Péclet number *Pe*; *Pe* ≫ 1 where advection dominates, and *Pe*≪ 1 where diffusion dominates. Across the brain, *Pe* varies by several orders of magnitude, with some regions dominated by advection and others dominated by diffusion (Figure 4). Diffusion dominates in most of the brain, but advection dominates in many regions where the velocity is large, e.g., near the olfactory bulb, cisterna magna, and Circle of Willis (Fig. 4A-C). Since *Pe* is proportional to the velocity, it is not surprising that high-velocity regions often overlap high-*Pe* regions. That said, maps of ||**u**|| and *Pe* are not identical, and PDFs of *Pe* are less bimodal than PDFs of ||**u**||, so diffusion must also affect *Pe*. The PDF of *Pe* is similar for all five mice, suggesting that the distribution results from shared anatomical features.

**Figure 4.**
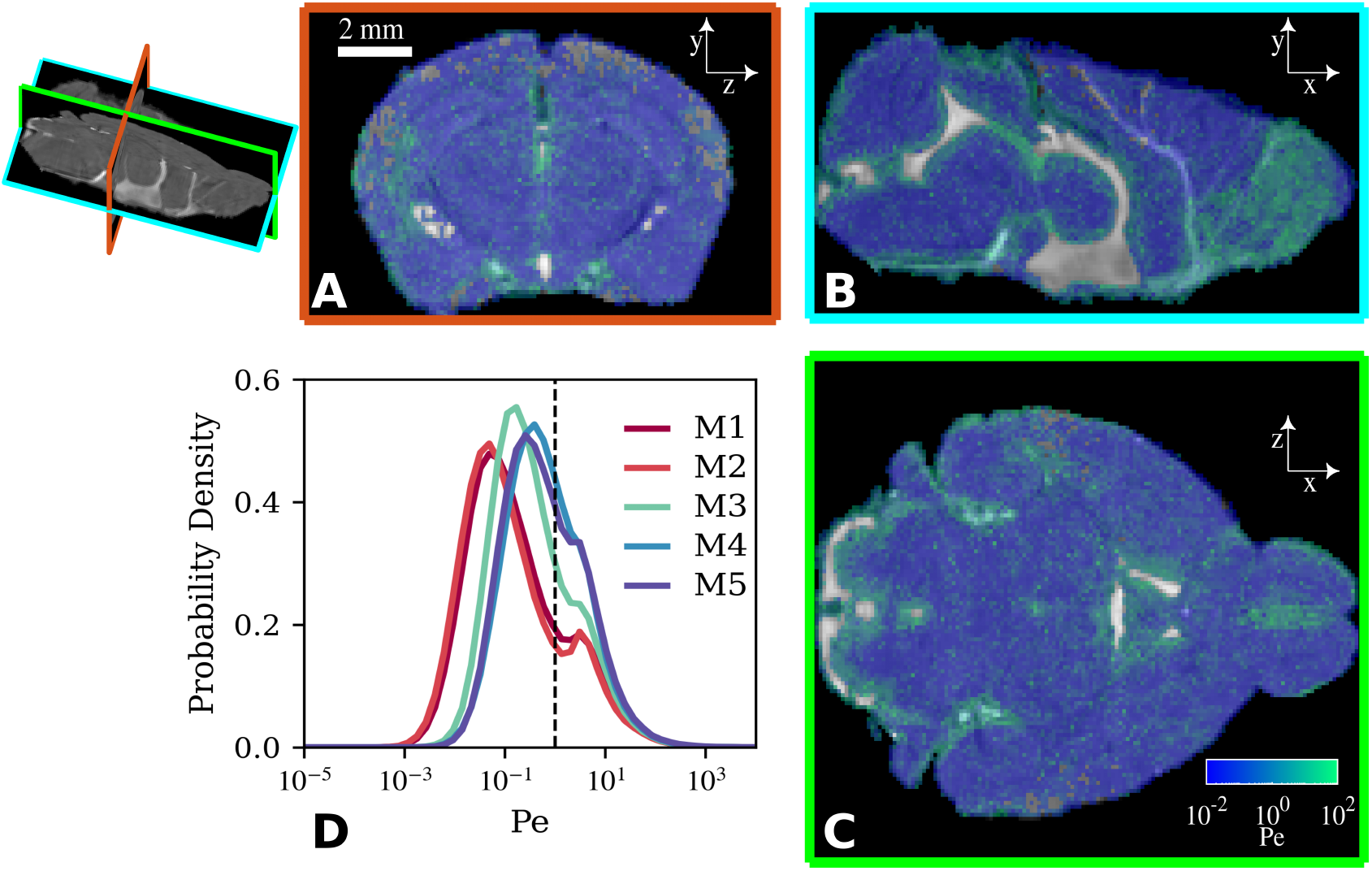
The local Péclet number varies spatially and is similar across mice. (A-C) Local Péclet number on mid-coronal (*z*-*y*), sagittal (*x*-*y*), and transverse (*x*-*z*) planes in Mouse 1. Solute is transported primarily by advection in green regions and diffusion in blue regions. A greyscale structural MRI image is visible in excluded regions. (D) The probability densities of local Péclet number throughout the brains of five mice (M1-M5).

#### 2.5. MR-AIV maps permeability and pressure

Maps of inferred permeability exhibit consistent spatial patterns across all five mice (Figure 5) and show, in more detail, the anatomical variation discussed above: high permeability around the olfactory bulb and perivascular spaces. Inferred permeability maps are similar across mice but differ from initial guesses, suggesting that spatial variations result from anatomical features. Inferred pressures are similarly consistent across mice. However, the absolute magnitudes of inferred permeability and pressure cannot be directly compared across mice due to the ill-posed nature of the underlying inverse problem.

**Figure 5.**
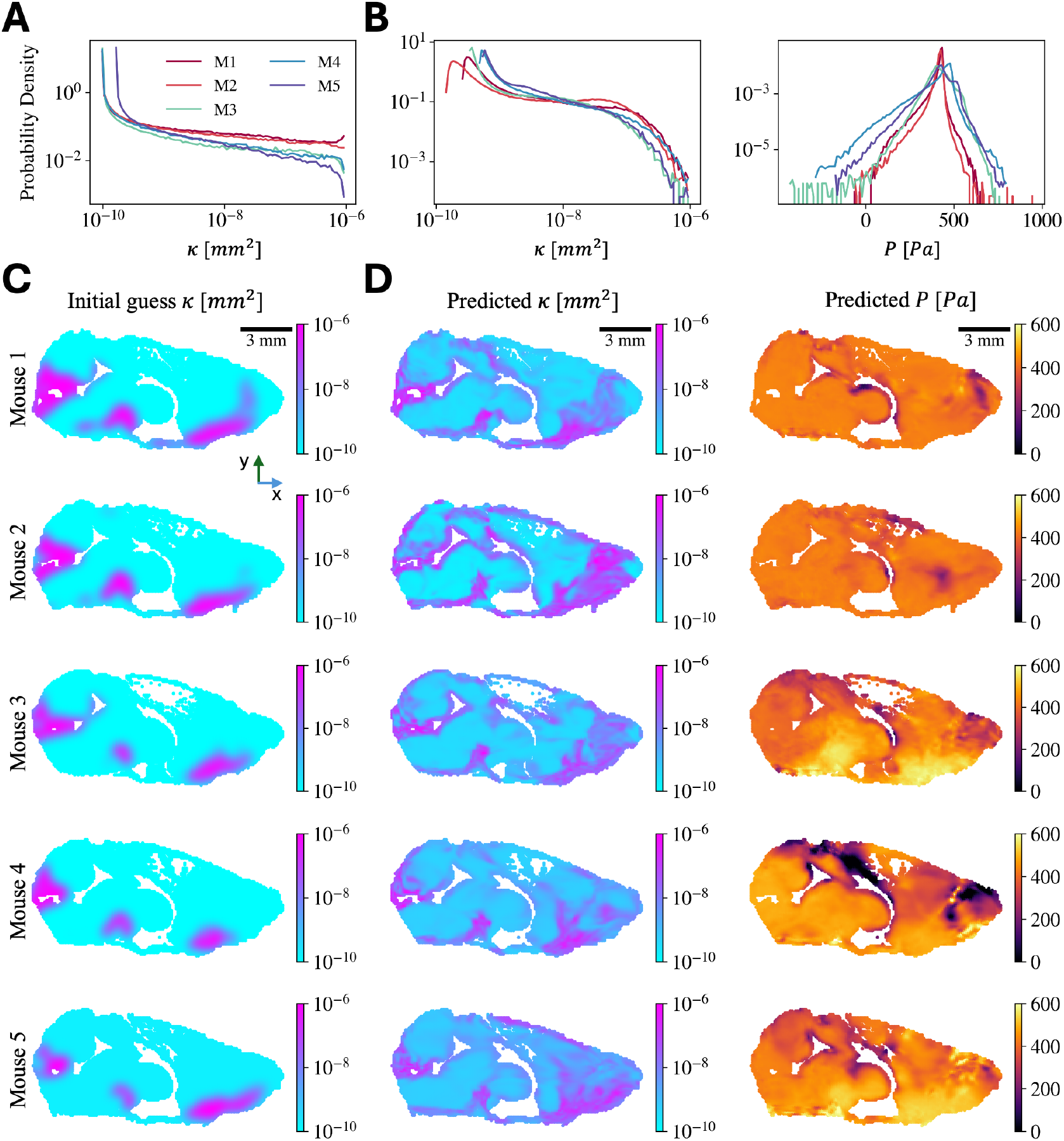
Estimated permeability and pressure. (A) Probability density functions of the initial permeability guess *κ* for each mouse (M1–M5). All distributions span more than four orders of magnitude, from 10^−10^ to 10^−6^ mm^2^, with little variability across mice. (B) Probability density functions of the estimated permeability *κ* and pressure *p* after training. (C) Initial permeability guess for each mouse (M1–M3). Whereas the overall dynamic range is consistent, spatial distributions differ across mice. (D) Predicted permeability *κ* and pressure *P* are spatially coherent and consistent across five mice.

#### 2.6. MR-AIV eliminates noise, trains on data, then trains on physics

The MR-AIV framework that infers these flows begins with pre-processing, where initial estimates of permeability *K* and **u** are obtained, then an initialization step that concurrently denoises the concentration data and aims to match the initial estimates, and a training stage, in which the inferred fields are refined to satisfy the governing physical laws (see Methods). During the pre-processing stage we estimate **u** using front-tracking [4, 11, 12].

The permeability of brain tissue is unknown, with estimates spanning multiple orders of magnitude [40], so in MR-AIV, the model learns the permeability, given an initial estimate. That estimate is made by assuming that regions quickly reached by tracer have high permeability, and other regions have permeability 10^4^-fold lower, and by subsequently applying some smoothing (see Methods). During the first step of the initialization (see Figure 6(B)), we account for noise in experimental data by incorporating two neural networks, *NN*_*c*_ and *NN*_*σ*_, which learn the mean concentration field and its associated time-dependent standard deviation (i.e., noise), respectively, by minimizing the negative log-likelihood of the observed concentration data [41].

**Figure 6.**
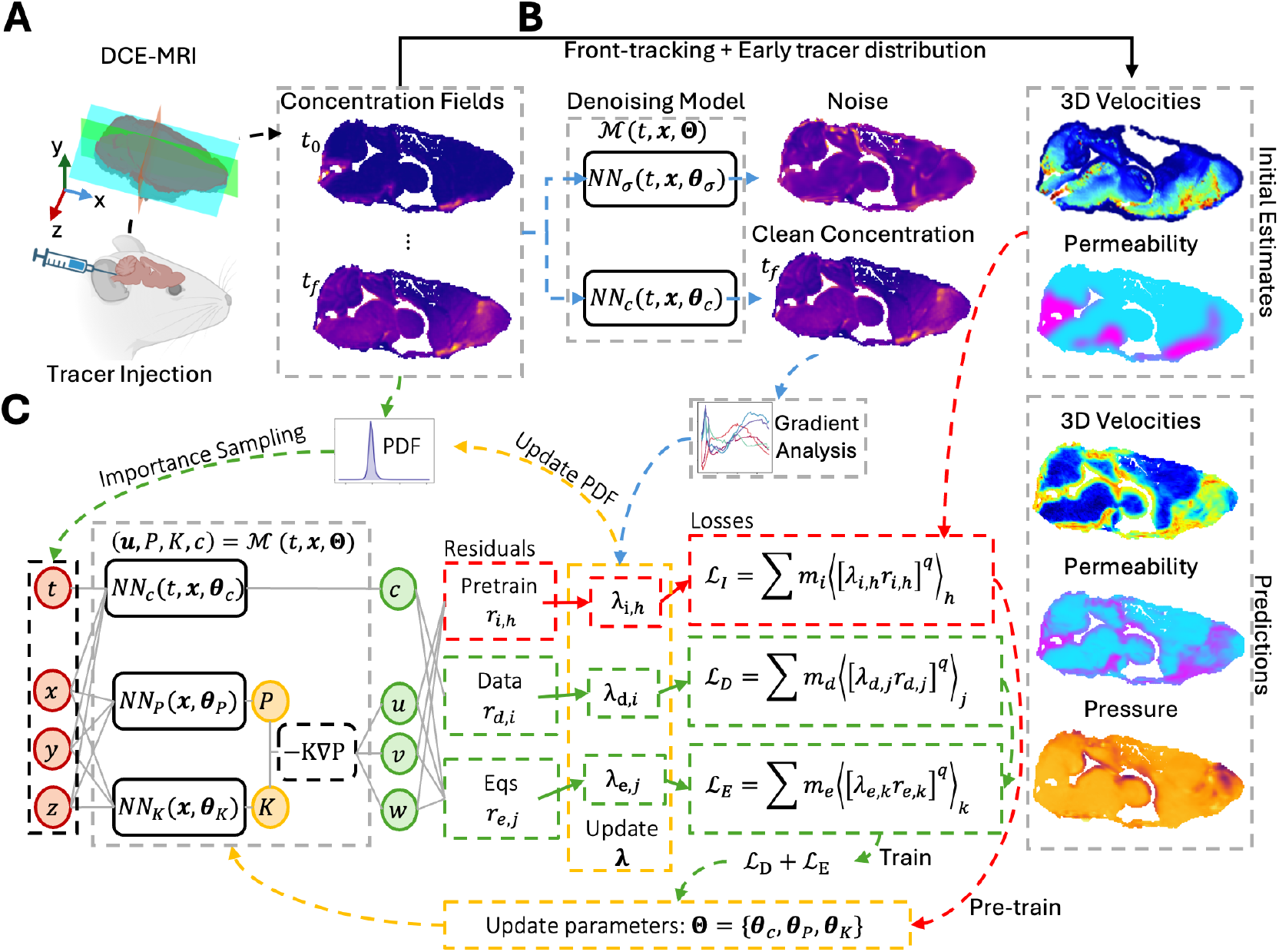
Magnetic Resonance Artificial Intelligence Velocimetry (MR-AIV) framework. MR-AIV infers 3D steady velocity **u**, permeability *K*, and pressure *P* fields from time-dependent concentration data. **(A)** Brain-wide concentration fields are obtained by injecting gadobutrol and tracking the tracer using DCE-MRI. **(B)** Initial estimates for **u** and *K* are derived from front tracking and early tracer distributions. A denoising module learns to separate experimental noise from the underlying signal (blue path). **(C)** The architecture enforces Darcy’s Law and steady-state conditions using independent networks for concentration, pressure, and permeability (*NN*_*c*_, *NN*_*p*_, *NN*_*k*_). The model is initialized by matching the initial estimates (red path), then refined by minimizing the residuals of the governing equations and concentration data (green path). Convergence is enhanced by time-dependent residual-based attention (TD-RBA), which uses weights (**˘**) based on residuals and clean concentration gradients to dynamically resample high-error regions, thereby improving convergence across the entire domain.

As shown in Figure 6(C), the MR-AIV architecture ℳ(**x, Θ**) with learnable parameters **Θ** = {***θ***_*c*_, ***θ***_*P*_, ***θ***_*K*_} consists of three neural networks: *NN*_*c*_(*t*, **x, *θ***_*c*_), which predicts the time-dependent concentration field *c* at any location **x** = (*x, y, z*); and *NN*_*P*_ (**x, *θ***_*P*_) and *NN*_*K*_(**x, *θ***_*K*_), which predict the steady-state pressure *P* (**x**) and permeability *K*(**x**), respectively. (We define the *x, y*, and *z* directions to be normal to the transverse, coronal, and mid-sagittal planes, respectively.) The velocity **u** = (*u, v, w*) can then be calculated via Darcy’s law: **u** = − *K*∇*P*.

During the second step of the initialization stage, the pressure and permeability networks are optimized using a data-driven approach that minimizes a loss function ℒ_*I*_, which penalizes the residuals *r*_*i*_ (i.e., point-wise errors) between the initial estimates and the predictions. Finally, during the training stage, predictions are refined by optimizing a combined loss function ℒ_*D*_ +ℒ_*E*_ that minimizes the residuals of measured concentration (*r*_*d*_) and the governing equations (*r*_*e*_) (see Figure 6(C)).

To promote uniform convergence during training, we introduce time-dependent residual-based attention (TD-RBA). This weighting approach improves previous residual-based attention strategies [37, 42–46] by computing attention weights *λ*_*i*_ based on the residual magnitudes and a physics-based time dependent scalar computed from the spatial gradients of the clean concentration field. TD-RBA are crucial for minimizing the multiscale residuals of governing equations and are used both as local multipliers to balance point-wise errors in the loss function and to define a residual-based probability density function (PDF) for resampling high-error regions. This resampling procedure improves predictions, allowing the model to focus training on regions with large residuals while maintaining stability across the full spatiotemporal domain. A comprehensive description of the MR-AIV architecture, sequential training strategy and the proposed (TD-RBA) is presented in Section 4 and Appendix A.

#### 2.7. MR-AIV is validated with synthetic data, and uncertainty with real data is quantified

To evaluate MR-AIV, we conducted a validation study on three synthetic data sets generated using finite element simulations that provided ground-truth velocity, pressure, and permeability fields (see Figure B.11 and B.12). Our validation confirmed that the framework successfully reconstructs concentration fields with high accuracy (relative *L*^2^ error *<* 2%, Figure B.10). For the more challenging task of velocity inference, performance depended on the complexity of the underlying permeability map. For the most complex “realistic” case, the average relative *L*^2^ error was 34% (see Table B.1).

However, as shown in Figure B.11, this error is not uniform; it is overwhelmingly concentrated in extremely low-velocity regions. This pattern is an expected consequence of the data: in slow-flow regions, minimal tracer is transported, resulting in a weak concentration signal that provides little information to constrain the velocity inference. Despite this inherent challenge of inferring a velocity field that spans more than four orders of magnitude, 43.1% of the domain still exhibited a relative error below 25% (see Figure B.11(A)). Furthermore, as shown in Figure B.11(B), the proposed framework successfully captures the bimodal velocity distribution, a key physical feature not retained by standard PIML models. In a simpler “smooth” case, the model performed optimally, with 88.5% of the domain having a relative error below 25%. Crucially, MR-AIV also successfully captured the correct spatial distributions of permeability and pressure (see Figure B.12), providing the first-ever estimates of these fields.

We then performed a robustness analysis on the initial permeability guess, which in turn assesses the epistemic uncertainty of our framework. In our model, the initial permeability guess effectively defines the parameter initialization for the subsequent velocity inference stage. We therefore quantified this uncertainty using an ensemble-of-models (EoM) approach, training multiple models with different initial permeability guesses. For the synthetic data, this analysis revealed a strong correlation between the predicted relative uncertainty and the true point-wise error (Fig. 7(A)), validating our uncertainty estimate as a reliable proxy for accuracy. When applied to the in vivo data, the uncertainty was highest in low-velocity regions, consistent with our synthetic findings (Fig. 7(B)). Despite the inherent difficulty of the problem, the uncertainty remained below 100% across 90% of the brain, providing informative, first-of-their-kind measurements of deep brain flow dynamics.

**Figure 7.**
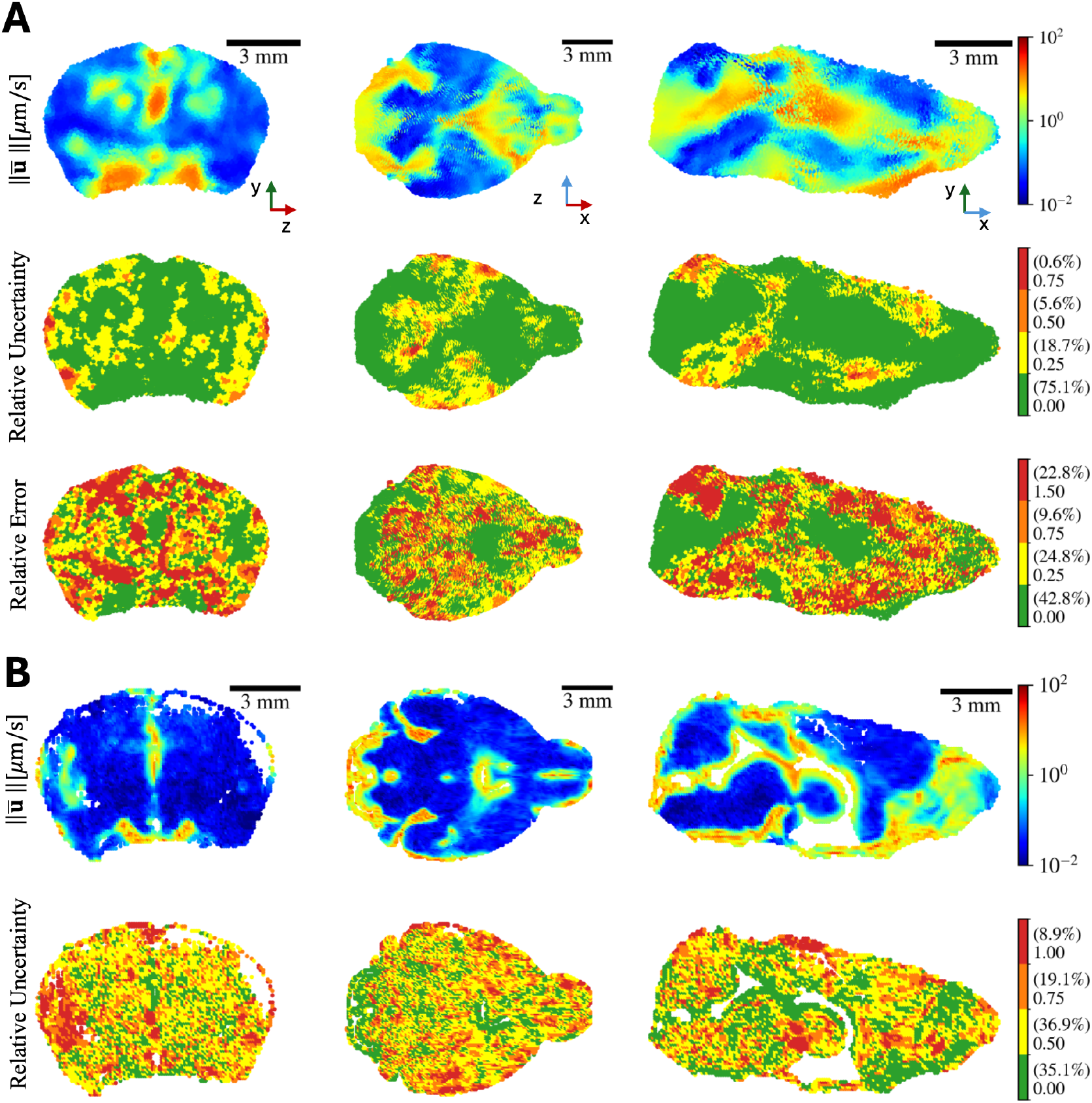
Uncertainty quantification of inferred velocity fields. **(A)** Uncertainty quantification for “realistic” synthetic data. The top row shows the velocity magnitude 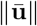 averaged across an ensemble of four models. The middle row shows the relative uncertainty 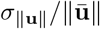, and the bottom row shows the point-wise relative error compared to ground truth. High uncertainty correlates with high error, and both are concentrated in low-velocity regions. The average relative *L*^2^ error for the ensemble is 34%. Parenthetical values indicate the percentage of total brain volume in each color. **(B)** Uncertainty quantification for Mouse 1. The top row shows the velocity magnitude averaged across the ensemble, and the bottom row shows the relative uncertainty. Consistent with the synthetic data, uncertainty is highest in low-velocity regions, yet remains below 100% across 90% of the domain.

## 3. Discussion

In summary, we introduce Magnetic Resonance Artificial Intelligence Velocimetry (MR-AIV), a specialized physics-informed machine learning framework designed to infer steady, brain-wide velocity, pressure, and permeability from time-dependent DCE-MRI concentration data.

MR-AIV can infer flows in the brain from sparse, noisy data without requiring velocity measurements, offering a generalizable and data-efficient tool for modelling transport in biological systems. We validated MR-AIV on synthetic data designed to mimic DCE-MRI measurements of tracer injected in the brains of mice and observed good agreement (see Figure B.11). An uncertainty quantification analysis demonstrated the robustness of the inferred velocities.

We used MR-AIV to quantify fluid flow and Péclet number throughout the entire brains of five wild-type mice. Typical velocity magnitudes, Péclet numbers, and anatomical variations of both were broadly similar across mice, suggesting the observed features are robust. PVSs and SASs harboured fast flow, while flow elsewhere was much slower, supporting the idea that PVSs and SASs serve a distinct role as highspeed pathways [47]. That distinct role is further supported by maps of the Péclet number: in small regions around PVSs, ventricles, and the olfactory bulb, *Pe* ≫ 1, implying advective solute transport, but elsewhere, *Pe* ≪ 1, implying diffusive solute transport. The finding of rapid, advection-dominated transport in the olfactory bulb is particularly significant, as this area has been identified as a primary entry route for environmental neurotoxins, such as inhaled iron nanoparticles that contribute to sex-specific, neurodegenerative pathologies [48]. Local velocity is apparently determined more by variations of permeability, which spans four orders of magnitude and correlates strongly with velocity (see Figure C.13E), than by variations of pressure, which spans just one. Thus, the brain seems to regulate flow primarily through tissue properties, not pressure sources. Across mice, permeability was large near ventricles, the olfactory bulb, the cisterna magna, and the Circle of Willis (see Figures 3 and Appendix B.2.2). Pressure was low in the olfactory bulb and high near the Circle of Willis. Inferred pressures reached 600 Pa, lower than the typical cardiac pulse pressure and of the same order as numerical estimates [49]. Future measurements of pressure gradients could test these predictions. Interestingly, we repeatedly observed fast flows near the ventricle walls; future work should explore the possibility of substantial flow there.

For each mouse, the largest peak in the PDF of velocity magnitude occurred near 0.1 µm/s, corresponding to velocities common in most of the parenchyma. For comparison, Ray et al. [27] estimated interstitial flow velocities of 0.1 µm/s by incorporating the possibility of bulk flow into a transport model of real-time iontophoresis and comparing with experimental measurements in mice. Vinje et al. [25] used forward and inverse subject-specific modelling to show that tracer transport in humans could be explained by a combination of diffusion and interstitial flow velocities 0.017 to 0.15 µm/s. Chen et al. [23] estimated velocities of 0.1 µm/s in the brain parenchyma of rats using DCE-MRI and regularized optimal mass transport (rOMT), an inverse method based on minimizing tracer displacement [20–24]. Thus, MR-AIV infers parenchymal velocities consistent with several prior estimates and supports the idea of slow but important bulk flow of interstitial fluid [50].

For each mouse, another peak in the PDF of the velocity magnitude occurred near 3 µm/s, corresponding to velocities common in open regions and similar to the 2-5 µm/s previously estimated for penetrating and capillary PVSs and parenchyma [14] and the ∼ 20 µm/s measured in pial PVSs [6, 9, 51, 52]. The fact that PVSs are smaller than a voxel probably contributes to our velocities, which are necessarily averaged over a voxel, being comparatively slow. Chen et al. [23] estimated velocities of 0.2 µm/s in large surface PVSs, 15-fold slower than the ∼ 3 µm/s inferred by MR-AIV. The difference may be explained by the fact that Darcy’s law is used in MR-AIV but not in rOMT. In particular, by incorporating Darcy’s law, our framework captures the multiscale nature of brain fluid flow, distinguishing high-flow conduits from the parenchyma, a distinction that is less pronounced in methods that do not enforce this physical constraint.

When we made models without Darcy’s law, predicted velocities spanned a smaller range and were unimodal, failing to distinguish high-speed pathways from other regions (Fig. C.14C). Though Chen et al. did not report PDFs of velocities, the fact that they found velocities in pial PVSs exceeding those in porous regions by only a factor of two suggests a narrower distribution, consistent with what we observed when we did not use Darcy’s law. Another possible explanation is image resolution: compared to the 100-µm voxels used here, the 234-µm voxels of Chen et al. would introduce stronger partial-volume effects. Voxel volumes being 12.8-fold larger might explain the estimated velocities being 15-fold smaller. On the other hand, the largest PVSs are bigger in rats than in mice, partly alleviating partial volume effects.

MR-AIV inferences from synthetic data showed higher errors and uncertainties in regions with slow flow (see Figure B.11 and C.14). There, diffusion tends to flatten concentration gradients, while advection transports tracer only weakly, so the effective signal-to-noise ratio for inferring velocity from advection is low.

Additionally, little signal is available in slow regions because the tracer tends to appear there only near the end of an experiment or simulation. At each time during an experiment, at least 40% of the concentration data was discarded because its local magnitude or gradient was too small to be useful. Despite sometimes having relatively large uncertainty (∼ 100%) in the low velocity regions (see Figure 7), inferences from MR-AIV are nonetheless informative, considering the scarcity of published measurements of flow velocities in the deep brain. The ability to estimate uncertainty with MR-AIV allows systematic improvement in future work.

The synthetic data used for validation was made as realistic as possible (see Figure B.9), with the domain shape taken from Mouse 1, and the inlet and outlet pressure boundary conditions chosen to produce velocity magnitudes similar to those inferred from the real data (Figure 7). The resulting velocity map matched anatomical expectations, with high velocities in the olfactory bulb, near the cisterna magna, and near the Circle of Willis, and low velocities in the brain’s interior.

Additionally, we calculated the epistemic uncertainty arising from the initial guess for the permeability, using the same permeability maps we used for the in vivo data, and we saw similar trends in the uncertainty map. Synthetic data captured key features of in vivo data, like bimodal velocity distributions that span more than four orders of magnitude. Our synthetic data is freely available and can be used to benchmark alternatives to MR-AIV.

MR-AIV infers permeability, pressure, and velocity by assuming that the observed concentration is governed by the advection-diffusion equation, the continuity equation, and Darcy’s law. Combining these three equations yields two expressions (Eq. A.4) involving products of permeability and pressure (or their derivatives). However, except in products, permeability and pressure do not appear, so multiple combinations of the permeability and pressure can yield identical concentrations and velocities: scaling permeability by a constant factor and pressure by the inverse of that factor has no effect on velocity. Without additional constraints, the inferred pressure and permeability fields are not unique (though velocity is).

That said, the spatial variations of permeability and pressure, aside from the scaling factor, are fully captured by Eq. A.4. Consistent with that fact, MR-AIV models trained on synthetic data infer the spatial variations remarkably accurately.

In vivo, true permeability is not known, but permeabilities inferred from experiments were more similar across mice than to the initial permeability guess (Fig 5C and D), and four different initial guesses resulted in similar inferred permeability fields (Fig. C.15), further supporting the idea that inferred permeabilities are valid. Future measurements of the absolute permeability or pressure in even one location would eliminate ambiguity from Eq. A.4 and allow MR-AIV to make unique inferences.

We constructed an initial guess of the permeability purely from the tracer concentration soon after injection (see Methods). This data-driven approach is repeatable and avoids bias associated with extensive user input. However, in future work, better guesses might come by combining early measurements with anatomical knowledge. In our guess, the maximum permeability was set by calculating the equivalent resistance of an open, 6-µm cylindrical vessel. Many open structures are larger and presumably allow faster flows, but nearly all are also smaller than a voxel, so voxel-averaged velocities and effective permeabilities would be reduced, offsetting the large size. Because the amount of reduction is unknown, the maximum permeability is uncertain. For low-permeability regions, published estimates vary widely, from 1 × 10^−11^ to 4.50 × 10^−9^ mm^2^ [29, 53]. We chose 1 × 10^−10^ mm^2^, though again, the value is uncertain. Accordingly, we estimated the epistemic uncertainty arising from the initial permeability field guess since we consider that it is the largest source of uncertainty in our modelling. More accurate measurements of brain permeability would substantially reduce the uncertainty stemming from the initial permeability guess. That said, uniformly scaling the entire initial permeability guess by a constant factor, even by an order of magnitude, changes our results little because the initial guess is non-dimensionalised and scaled during calculations. As shown in equation A.6, a scaling factor would effectively redefine the characteristic permeability and reciprocally modify the characteristic pressure without changing the velocity.

We used the scaled SER as a proxy for tracer concentration, implying a linear relationship between the two quantities. Ratner et al. [20], using a similar imaging protocol, suggested that the relationship is nearly linear for SER less than 200%, which is true for *>* 99% of our individual measurements.

MR-AIV is limited to approximating velocities, permeabilities, and pressures as steady. The true quantities certainly vary over time, at least by small amounts. Future improvements might allow MR-AIV to infer temporal variations, if enough information can be gleaned from concentration fields. That said, all our synthetic data had truly steady velocities, permeabilities, and pressures. Additionally, our experiments were performed with unperturbed, anaesthetised mice whose brain activity presumably varied relatively little.

Considering the prevalence of DCE-MRI data, the fact that it is one of the few approaches for visualising in vivo fluid flows in humans, and the insight into the glymphatic system that this imaging modality has already provided, brain-wide velocity fields inferred by MR-AIV may unlock key questions about glymphatic flows in the future. Integrating these novel fluid dynamics maps with state-of-the-art anatomical resources, such as high-resolution, distortion-corrected brain atlases [39, 54], and correlating them with other advanced modalities that probe tissue microstructure [55], represents a promising direction for creating a more complete biophysical model of the brain.

## 4. Methods

### 4.1. Imaging data collection and pre-processing

Five 11-13-week-old male C57BL/6 wild-type mice were used for this experiment. The MRI data were collected in a 9.4 T preclinical scanner (BioSpec 94/30 USR, Paravision 6.0.1 software, Bruker BioSpin, Ettlingen, Germany) equipped with a 1 H cryogenically-cooled quadrature-resonator Tx/Rx coil (CryoProbe, Bruker) and a 240 mT/m gradient coil (BGA-12S, Bruker) at the Preclinical MRI Core Facility, University of Copenhagen. During the imaging, all the mice in the MRI scanner were placed in the prone position and anaesthetized with the intraperitoneal injection of the mixture of ketamine and dexmedetomidine (K/Dex: 75/1 mg/kg). The body temperature was maintained at 37 ± 1^°^C with a MR-compatible and thermostatically controlled waterbed and monitored by a remote monitoring system (SA Instruments, NY, USA), together with the respiratory rate. A stereotactic holder with ear bars was used to minimize the head movement during imaging. To investigate fluid flow dynamics, a contrast agent was injected intracisternally as previously described [56, 57]. Briefly, a 30G copper needle (outer diameter 0.32 mm; Nippon Tokushukan, Mfg, Tokyo, Japan) attached to a PE10 tubing was inserted into the cisterna magna. A T2-weighted structural image was conducted using 3D constructive interference steady-state (3D-CISS). Each 3D-CISS image was calculated as a maximum intensity projection of 4 realigned 3D TrueFISP volumes with 4 orthogonal phase encoding directions (TR/TE 3.9/1.95 ms, Nex 1, FA 50^°^, FOV 19.2 × 12.8 × 12.8 mm, Matrix 192 × 128 × 128). For dynamic contrast-enhanced magnetic resonance imaging (DCE-MRI), T1-weighted imaging was acquired using a 3D-FISP sequence (TR/TE 4/2 ms, Nex 1, FA 15^°^, FOV 19.2 × 12.8 × 12.8 mm, Matrix 192 × 128 × 128). First, three baseline DCE-MRI scans (3 min) were acquired. T1-enhancing contrast agent gadobutrol (15 mM; Gadovist, Bayer Pharma AG, Leverkusen, Germany) was infused into the cisterna magna. Over a 10-minute period, 10 µL of gadobutrol was infused at a rate of 1 µL/min utilizing a 100 µL Hamilton syringe mounted on a motorized pump. The image spatial resolution was 100 µm isotropic (0.001 mm^3^) with one scan every 60 seconds. After the three baseline scans, the follow-up scans continued over 90 minutes. Further image processing pipelines were applied, including motion correction, bias field correction, and spatial co-registration between 3D-CISS and DCE-MRI images. To normalize the image in each time series, their voxel intensities were subjected to Gaussian normalization using the first 3D-FISP volume. The resulting images were smoothed with a 3 × 3 × 3-voxel kernel of [0.2, 1, 0.2] weights along each axis, to reduce the influence of possible artefacts after the automatic registration and subtracting the baseline volume. Voxel-based percentage enhancement of contrast from baseline (signal enhancement ratio, SER) was calculated as equation of *SER* = (*S*_*t*_ − *S*_0_)*/S*_0_ × 100 (Figure C.13A).

To avoid using a source term in our governing equations, we excluded the concentration data in the cisterna magna. We also excluded the ventricles, where CSF is produced, regions outside of the brain, and regions that the tracer did not reach during the 93-minute scan duration (defined as regions where the change in SER through all scans was less than three times the change in SER during the baseline portion of the scan). The excluded regions are shown on a mid-sagittal slice in Figure C.13B. Accordingly, there are no concentration, velocity, or permeability predictions in the excluded regions.

### 4.2. Initial velocity estimate from front tracking

We used front tracking to provide an initial estimate of the velocity field, which is used to initialize the neural networks. In front tracking, fronts, or regions of constant concentration, are tracked over time and space. Front tracking neglects diffusion and thus has low accuracy in regions where diffusion plays a substantial role in transport. The front tracking algorithm we employ is inspired by one described by Nevins and Kelley [58, 59] but has been adapted for three dimensions and does not fit curves to the fronts. We combine sparse velocity information from many different fronts corresponding to various concentrations and assume the flow is steady, thereby obtaining estimates of the velocity at nearly every location tracer reaches in the brain. Figure C.13D shows example velocity estimates from front tracking.

### 4.3. Material property estimates

The material properties of CSF and ISF are similar to those of water, so we used the density and viscosity of water at 37^°^C (density *ρ* = 993 kg/m^3^, viscosity *µ* = 6.95 × 10^−4^ Pa · s). We used *D* = 2.4 × 10^−4^ mm^2^/s for the diffusion coefficient of gadobutrol. The free diffusion coefficient for Gd-DPTA, which has a similar molecular weight as gadobutrol (550 Da compared to 604 Da) is *D*_free_ = 3.8 × 10^−4^ mm^2^/s [13], but since most of the brain tissue is porous, the effective diffusion coefficient in the porous regions is *D*_eff_ = 1.48 × 10^−4^ mm^2^/s (where *D*_eff_ = *D*_free_*/λ*^2^, assuming the tortuosity is *λ* = 1.6, which has been repeatedly measured for ECS). This also agrees with estimates of the diffusion coefficient from Ringstad et al. [60]. Since the diffusion coefficient number only varies by a factor of 2.5 between open and porous regions, we used the constant value *D* = 2.4 × 10^−4^ mm^2^/s everywhere in the domain.

Identifying open spaces within the brain, particularly those that are smaller than an MRI voxel (e.g., PVSs), is not trivial, and most voxels containing “open” spaces also contain porous tissue. Therefore, we modelled the entire brain as a porous medium with spatially varying permeability to be learned by the neural network. We provided an initial estimate of the permeability map by assuming that regions that tracer reached (defined as having a SER greater than 150%) quickly (within 16 minutes) had high permeability (1 × 10^−6^ mm^2^, the effective permeability of an open cylindrical channel with a diameter of 6 µm), and all other regions had low permeability (1 × 10^−10^ mm^2^). Though high-permeability voxels are likely to contain open spaces larger than 6 µm, because they also contain porous medium, we did not set the effective permeability higher. The most accurate permeability would be a weighted average across the voxel’s volume, but no methods yet exist to provide the necessary information. We smoothed the binary permeability map using a 3D Gaussian filter with a standard deviation of one or three voxels for the “sharp” and “smooth” cases, respectively. Since the high-permeability regions are small, smoothing sometimes lowers their maximum permeability; we then scaled *κ* to return its maximum to 1 × 10^−6^ mm^2^. The velocities shown in Figures 1 and 3 come from the “smooth” initial permeability maps shown in Figure 5.

### 4.4. Underlying Physical Laws

We modeled tracer transport using the advection-diffusion (Eq. A.1) and continuity (Eq. A.2) equations, assuming an incompressible flow. However, inferring the three-dimensional velocity field from concentration data alone is a severely ill-posed inverse problem; with only two governing equations for three unknown velocity components and sparse concentration data, the system is underdetermined.

To constrain the problem, we modeled the brain as a porous medium, which allows fluid flow to be governed by Darcy’s law (Eq. A.3). Incorporating Darcy’s law is critical for two reasons. First, it closes the system by expressing the velocity field in terms of permeability and pressure, reducing the number of unknown variables to be solved. Second, the permeability field acts as a learned spatial map, enabling the model to capture the sharp transitions between high- and low-velocity regions that span several orders of magnitude. The analysis was performed over a 90-minute period following an initial 5-minute injection phase. The full, non-dimensionalized governing equations used in our framework are detailed in the Appendix A.1.

### 4.4.1. Local Péclet number

The local Péclet number *Pe* is a dimensionless parameter that describes the relative contributions of advection and diffusion to mass transport, defined as

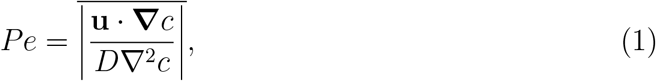

where the operator 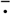 denotes an average over time.

### 4.5 Magnetic-Resonance Artificial Intelligence Velocimetry (MR-AIV)

MR-AIV is a scientific machine learning framework inspired by AIV [9, 10, 36] and AIVT [37], designed to infer continuous and differentiable steady-state velocity fields ***u*** = (*u, v, w*) from time-dependent, sparse concentration measurements *c*. For robustness and stability [61], MR-AIV approximates the solutions of the governing equations A.3, A.2, and A.1 in their non-dimensional form as described in Appendix A.1:

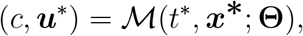

where **Θ** represents the trainable parameters of the model, (*t*^*^, ***x***^*^) = (*t*^*^, *x*^*^, *y*^*^, *z*^*^) denotes the non-dimensional temporal and spatial inputs, and *c* and ***u***^*^ = (*u*^*^, *v*^*^, *w*^*^) are the non-dimensional concentration and velocity fields (see Appendix A.1). As shown in Figure 6(C), the proposed model *ℳ* (·) is composed of three independent neural networks: *NN*_*c*_(·), *NN*_*K*_(·), and *NN*_*P*_ (·), which respectively approximate the non-dimensional concentration (*c*), pressure (*P* ^*^), and permeability (*K*^*^) fields:

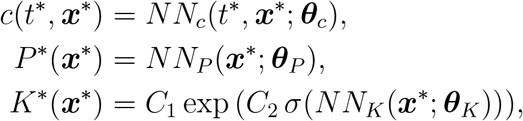

where ***θ***_*c*_, ***θ***_*P*_, and ***θ***_*K*_ denote the parameters of each neural network. The sigmoid function *σ* : R → [0, 1] is used in combination with the constants *C*_1_ = *K*_max_ and 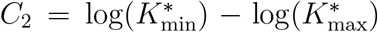 to ensure that permeability remains bounded between 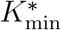 and 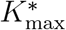. The non-dimensional velocity field ***u***^*^ is then recovered from *K* and *P* ^*^ via ***u***^*^ = − *K*^*^ ∇ *P* ^*^, thereby strictly enforcing the non-dimensional Darcy’s law. This formulation also inherently satisfies the steady-state assumption, as both *K*^*^ and *P* depend solely on the spatial coordinates ***x***^*^. All MR-AIV subnetworks (*NN*_*c*_(·), *NN*_*σ*_(·), *NN*_*K*_(·), and *NN*_*P*_ (·)) in this study use weight normalization [62] to speed up convergence and adaptive residual connections to avoid vanishing gradients [63]. Additionally, the inputs of all models use a fifth-degree polynomial feature expansion to improve the model representation capabilities [44]. The remaining details about the architecture and implementation are described in the Appendix D. Once the model is trained, we obtain the corresponding dimensional fields as follows:

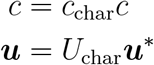

where *c*_char_ and *U*_char_ are the corresponding characteristic dimensions as specified in the Appendix A.1.

#### 4.5.1. Training

Since MR-AIV encodes Darcy’s law directly into its architecture, training a model involves optimizing two objective functions: the advection-diffusion equation and continuity as a PDE constraint, and the data fidelity term. However, this process requires inferring the velocity field from experimental concentration data, which is inherently noisy. Additionally, the model must learn the permeability field, which spans four orders of magnitude. Without proper initialization, these factors can lead the model to become trapped in poor local minima. To mitigate this issue, we divide the training process into two stages: initialization and full training.

During the initialization stage, we initialize the network parameters **Θ** = {***θ***_*c*_, ***θ***_*K*_, ***θ***_*P*_} using the experimental concentration data together with the initial estimates of pressure and permeability. This stage is further divided into two main steps: first, learning the experimental concentration data and its associated noise structure by minimizing the negative log-likelihood loss ; and second, fitting the initial permeability estimates, followed by refining the pressure field to match the initial velocity estimates (see Appendix A.2).

During the training stage, we refine the initial field estimates by incorporating the physical constraints, ensuring that the model produces physically consistent solutions that also match the observed concentration data. Towards this end, we minimize a combined loss function that penalizes both data mismatch and equation residuals. Each loss term is defined as:

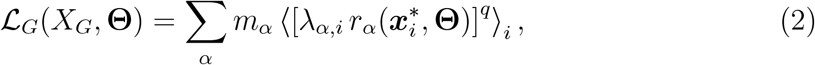

where 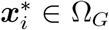 and *G* ∈ {*D, E*} indicates the loss group: data (*ℒ*_*D*_) or physics-based equation constraints (*ℒ*_*E*_). The operator ⟨·⟩_*i*_ denotes the mean over the training points 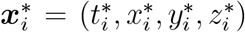 sampled from the subset *X*_*G*_ ⊂ Ω_*G*_. Sampling is performed iteratively using a probability distribution *p*_*G,α*_, defined using a residual-based-attention resampling (RBAR) [37] strategy as described in Appendix A.2. The exponent *q >* 0 controls the smoothness of the loss function, allowing for a transition between *L*^2^ and *L*^1^ norms (with *q* = 2 and *q* = 1, respectively) during training.

Each loss group *G* targets specific physical quantities, indexed by *α*. For the physics-based loss (*G* = *E*), we enforce the advection-diffusion equation and conservation of mass in their residual form, corresponding to *α* = {AD, CM}. For the data loss (*G* = *D*), we constrain only the concentration field, with *α* = {*c*}. The specific details of each loss subterm are described in Appendix A.2.

The residual 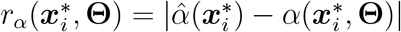 quantifies the discrepancy between the predicted value 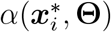 and the target value 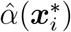 (e.g., experimental observation or zero for the PDEs). Finally, residual-based-attention (RBA) weights [43] *λ*_*α,i*_ are used as local multipliers to control the point-wise contribution of each residual, while global weights *m*_*α*_ balance the relative importance of each subcomponent *α* across the loss groups. The specific formulation for the RBA weights is described in the Appendix A.2.

##### Time-dependent Residual-Based Attention with Resampling (TD-RBA)

A key challenge in this problem is the vast dynamic range of the concentration field, which causes the physical residuals in the loss function to vary by orders of magnitude over time. To ensure stable training, we developed Time-dependent Residual-Based Attention (TD-RBA), a novel optimization method that acts as a point-wise adaptive learning rate, guided directly by the physics of the advection-diffusion equation. This is achieved by introducing a time-dependent weight for the advection-diffusion loss, *λ*_*AD,i*_(*t*_*i*_) = *λ*_*AD,i*_*/C*(*t*_*i*_), where *λ*_*AD,i*_ is a standard RBA weight. The main contribution of this study is the time-dependent scaling factor, *C*(*t*_*i*_), which normalizes the residuals at each time point and is defined by the maximum magnitude of the advection-diffusion equation’s components:

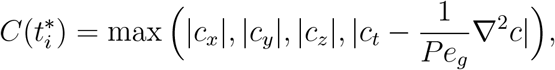

where all gradients are evaluated across the spatial domain at time *t*^*^*i*. This approach ensures all phases of tracer transport contribute meaningfully to the solution. The full formulation, including the underlying residual-based attention with resampling (RBA-R), is provided in the Appendix A.3.

### 4.6. Computational Fluid Dynamics Validation

To evaluate the performance of MR-AIV, we conduct a comprehensive validation using high-fidelity synthetic data sets generated via the finite element method. Each simulation consists of approximately 2 million tetrahedral elements (see Figure B.9(A)). Full details of the numerical setup are provided in the Appendix B.

We evaluate MR-AIV using three permeability maps to solve Darcy’s law, each yielding a distinct velocity field. To match the scale of real data, low and high permeability regions are set to 10^−6^, mm^2^ and 10^−10^, mm^2^, respectively. While all maps are predominantly binary, they differ in the sharpness and complexity of the transitions between regions. The “smooth” map contains gradual transitions, the “sharp” map features abrupt interfaces, and the “realistic” map combines sharp transitions with a more intricate spatial layout (see Figure B.9(B)). Once the velocity fields are obtained, we solve the advection-diffusion equation to compute the corresponding concentration fields.

We use 50% of the concentration data to train our MR-AIV model following the same strategy as for the real data. Then we evaluate performance using the relative *L*^2^ error quantified as:

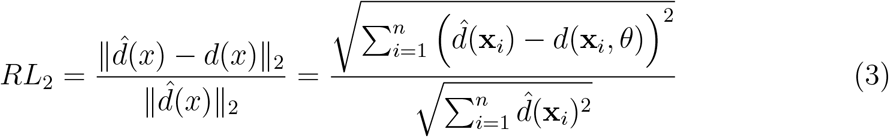

where *d*(*x*_*i*_) is the predicted value (i.e., concentration, velocity, permeability, or pressure) at point *x*_*i*_.

The model successfully reconstructed the synthetic concentration fields (Figure B.10), maintaining a relative *L*^2^ error below 2%. For the velocity magnitude inference (Figure B.11), relative errors were 36% as detailed in Table B.1. These inference errors were primarily concentrated in regions of extremely low velocity, a finding corroborated by the uncertainty quantification analysis as shown in Figures 7 and C.14.

Finally, the results for the estimated pressure and permeability are shown in Figure B.12. Notably, the predicted fields exhibit strong agreement with the reference fields, preserving key structural features and regional contrasts. However, while the spatial trends are well recovered, the absolute magnitudes of the predicted fields deviate from the ground truth due to the ill-posedness of the problem, as detailed further in Appendix A.3 and their corresponding uncertainty quantification (Figures C.15).

## Acknowledgements

We are grateful for many insightful conversations with Maiken Nedergaard.

## Author contributions

1. **Conceptualization:** G.E.K., D.H.K., J.D.T.
2. **Methodology:** J.D.T., K.A.S.B., Z.W.
3. **Software:** J.D.T., Y.G.
4. **Validation:** J.D.T., Y.G.
5. **Formal analysis:** J.D.T., M.V.
6. **Investigation:** J.D.T., Y.G., Y.M., K.A.S.B.
7. **Resources:** Y.M.
8. **Data curation:** K.A.S.B., Y.G., Y.M.
9. **Writing – original draft:** J.D.T., K.A.S.B., D.H.K.
10. **Writing – review & editing:** J.D.T., Y.G., Z.W., M.V., Y.M., G.E.K., K.A.S.B., D.H.K.
11. **Visualization:** J.D.T., M.V., K.A.S.B.
12. **Supervision:** G.E.K., K.A.S.B., D.H.K.
13. **Project administration:** G.E.K., D.H.K.
14. **Funding acquisition:** G.E.K., D.H.K.

## Appendix A. Materials and Methods

### Appendix A.1. Underlying Physical Laws

Tracer transport was assumed to follow the advection-diffusion equation with constant diffusivity *D*:

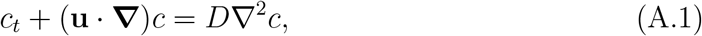

where *c* denotes the tracer concentration. The tracer evolves over the spatiotemporal domain Ω = Ω_**x**_ × Ω_*t*_ ⊂ ℝ^3+1^, with Ω_*t*_ = {*t* ∈ ℝ | *t*_0_ ≤ *t* ≤ *t*_*f*_}. Here, *t*_*f*_ = 90 minutes corresponds to the final acquisition time, and *t*_0_ = 5 minutes is selected to exclude the initial injection phase, since the specific source term in the advection-diffusion equation was unknown. We assumed that the flow was incompressible and imposed conservation of mass in the form

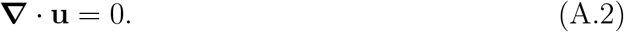

We constructed MR-AIV models using only the advection-diffusion and mass equations, setting out to infer the velocity **u**. Those models performed poorly, producing concentration and velocity fields that did not match the synthetic data well. We explain the poor performance with the fact that there were three unknown velocity components (and concentration, which is known only partially), but only two equations; the system may not have been closed.

MR-AIV models performed much better when we further asserted that brain tissue was a porous medium where fluid flow is governed by Darcy’s law:

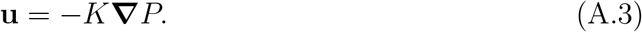

Here, **u** is the velocity field, *K* is the hydraulic permeability, and *P* is the pressure. Incidentally, *K* = *κ/µ*, where *κ* is the intrinsic permeability and *µ* is the fluid viscosity. The velocity was assumed to be steady within the spatial domain Ω_**x**_ = {**x** ∈ R^3^ | **x** ∈ *G*_*m*_}, where *G*_*m*_ denotes the set of spatial locations defining the brain geometry of each individual mouse *m*. Using Eq. A.3 to eliminate **u** from Eqs. A.1 and A.2 yields

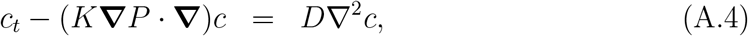

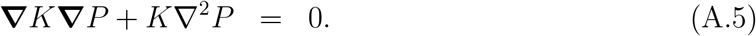

Since *D* is a known constant and *c* comes from measurements or simulations, this system of two equations has just two unknowns, *K* and *P*, and is therefore closed. We believe that its closure was essential for making accurate machine learning models. Once *K* and *P* have been inferred, **u** can be calculated using Eq. A.3.

In our physics-informed machine learning models, these equations are enforced in their non-dimensional forms, as described in detail in the Appendix A.1.

#### Non-dimensionalization

The concentration spans over four orders of magnitude, the permeability over four, and the velocity over five. To enable stable training and prevent numerical instabilities, it is therefore essential to non-dimensionalise these quantities. We define the following characteristic scales:

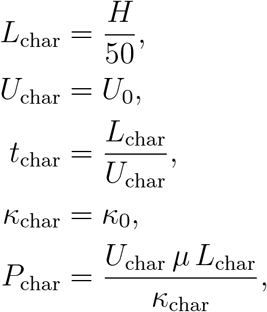

where *H* = 17 mm approximates the maximum length of the brain of mouse 1, and *U*_0_ = 0.1 mm*/*min corresponds to the expected velocity obtained from our front-tracking analysis. The characteristic permeability *κ*_0_ = 10^−8^ mm^2^ is set to one hundredth of the maximum permeability guess.

The physical quantities are then non-dimensionalised as

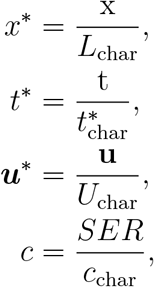

where t denotes the dimensional time, x, y, z are the dimensional spatial coordinates, and **u** = (u, v, w) represents the corresponding dimensional velocity components. The characteristic concentration 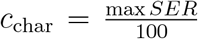is chosen such that the non-dimensional concentration falls within *c* ∈ [0, 100], and is therefore mouse-independent.

Finally, the non-dimensionalised pressure *P* ^*^ and permeability *K*^*^ are defined as:

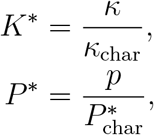

where *κ* and *p* denote the dimensional permeability and pressure field, respectively. Using these non-dimensional quantities, we rewrite the governing equations as:

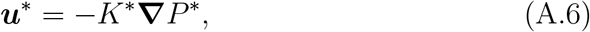

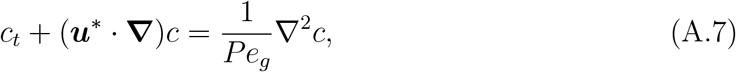

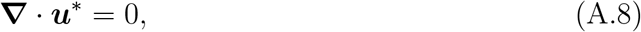

where *Pe*_*g*_ = *U*_char_*L*_char_*/D* = 2.361 is the global Péclet number, with *D* = 2.4 × 10^−4^ mm^2^*/*s denoting the diffusivity.

#### Equation Residuals

To enforce the governing equations within the PIML framework, we write them in residual form. The residuals used to constrain the non-dimensional advection-diffusion equation and the conservation of mass are defined as:

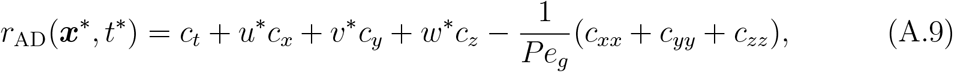

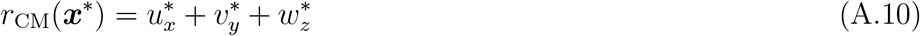

where the subscript denotes differentiation with respect to the corresponding variable (e.g.,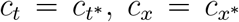). All spatial and temporal derivatives are computed using automatic differentiation. Note that Darcy’s law is not included as a residual term, as it is directly encoded into the model architecture.

### Appendix A.2. MR-AIV framework

#### Appendix A.2.1. Denoising Module via Negative Log-likelihood

Following [10], we use the negative log-likelihood (NLL) in PIML to explicitly model the aleatoric uncertainty (uncertainty due to noise) in the concentration data. Specifically, we assume that the observed concentration field *c*_obs_ can be decomposed into a noise-free prediction 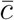 and additive Gaussian noise *ϵ* with zero mean and with spatially and temporally varying standard deviation *σ*_*c*_, such that

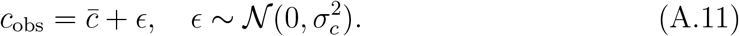

In this study, we model both the mean concentration field 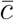and the corresponding standard deviation *σ*_*c*_ using two separate neural networks, with parameters ***θ***_*c*_ and ***θ***_*σ*_, respectively. The mean field 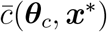 provides a denoised estimate of the concentration, while *σ*_*c*_(***θ***_*σ*_, ***x***^*^) captures the spatially and temporally varying uncertainty, where ***x***^*^ = (*t*^*^, *x*^*^, *y*^*^, *z*^*^) are the non-dimensional inputs.

To ensure numerical stability and physical plausibility [10], we bound the inferred standard deviation as

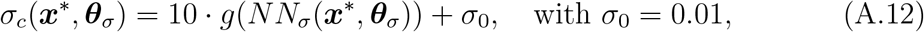

where *g* : ℝ → (0, 1) is the sigmoid function:

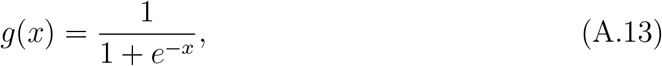

which ensures that *σ*_*c*_(***x***) ∈ (*σ*_0_, 10 + *σ*_0_) and prevents the network from predicting arbitrarily small or negative uncertainties. The negative log-likelihood (NLL) loss is then defined as:

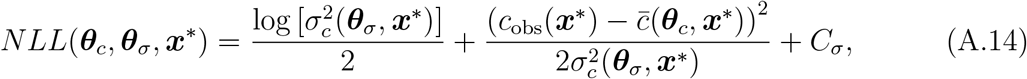

where the constant 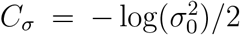 normalizes the metric such that the NLL is zero when the residual vanishes and *σ*_*c*_ = *σ*_0_. The denoising loss *ℒ*_*N*_ learns both the noise structure and the clean concentration field by penalizing the mismatch between the reconstructed mean concentration 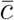 and the experimental observations *c*_obs_. It is explicitly defined as

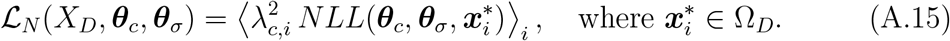

Here, ⟨·⟩_*i*_ denotes the mean operator over the batch *X*_*D*_ selected from the data domain Ω_*D*_. The residual-based attention (RBA) weights *λ*_*c,i*_ adjust the point-wise contribution of the residuals, enabling uniform convergence.

#### Appendix A.2.2. Sequential Training

##### Initialization

During this stage, we initialize the network parameters **Θ** using concentration measurements along with the initial estimates of pressure and permeability. This stage is further divided into two steps: first, fitting the initial concentration and permeability estimates, and second, refining the pressure to match the initial velocity estimates.

**Step 0:** We initialize the parameters ***θ***_*K*_ of the permeability network *NN*_*K*_ by training the model to match the initial permeability estimates. This is done by minimizing the first initialization loss 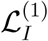, defined as:

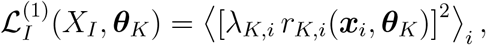

where 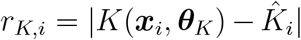 is the residual (point-wise error) between the prediction *K*(***x***_*i*_, ***θ***_*K*_) and the initial permeability guess 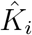 at location ***x***_*i*_ = (*x*_*i*_, *y*_*i*_, *z*_*i*_), and *λ*_*K,i*_ is the corresponding local weight, which balances the contribution of each training point to promote uniform convergence. Also, since the model would learn two regions that are clearly defined, we further split the initialization into two sub losses for the high and low permeability regions, respectively. We initialize the permeability network for *N*_*s*_0 = 500000 ADAMw [64] iterations.

**Step 1:** We initialize the parameters of the concentration network ***θ***_*c*_ using our denoising module, which simultaneously recovers a clean concentration field and estimates the noise distribution. During this step, the denoising loss *ℒ*_*N*_ is introduced to enable the learning of both the noise structure and the clean concentration by minimizing the negative log-likelihood of the experimental concentration data (equation A.14).

Concurrently, since the permeability was initialized in the previous step, we fix ***θ***_*K*_ and proceed to initialize the parameters of the pressure network ***θ***_*P*_ by fitting the initial velocity estimates obtained from the front-tracking analysis. Constraining the velocity in this way also implicitly constrains the pressure network through Darcy’s law, since the pressure gradients govern the velocity field. The corresponding initialization loss is defined as:

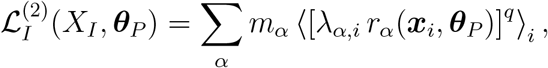

where *α* = {*u, v, w*} indexes the velocity components, *r*_*α*_(***x***_*i*_, ***θ***_*P*_) is the residual between the inferred and initial velocity estimates for each component, *m*_*α*_ is the global weight balancing the contribution of each sub-term, and *λ*_*α,i*_ is the corresponding local weight. We initialize the concentration and pressure for *N*_*s*1_ = 100000 ADAMw [64] iterations.

##### Filtering

Once we obtain the concentration field from the denoising module, the relatively high spatial resolution allows us to approximate the true gradients using automatic differentiation. Figure A.8(A) shows the time evolution of the concentration gradients *c*_*x*_, *c*_*y*_, *c*_*z*_, and 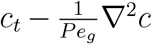. The results indicate that the maximum values of these gradients exhibit high fluctuations, which may be associated with measurement noise. Moreover, the gradient magnitudes vary substantially across times, spanning several orders of magnitude. This variability highlights a fundamental challenge for stable velocity inference. Therefore, under the assumption that the concentration gradients *c*_*x*_, *c*_*y*_, *c*_*z*_, and 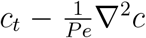 follow a log-normal distribution, we filter the data by discarding points that fall outside three standard deviations from the mean in the log-scaled domain. The mean and standard deviation of the log-scaled gradients are computed as follows:

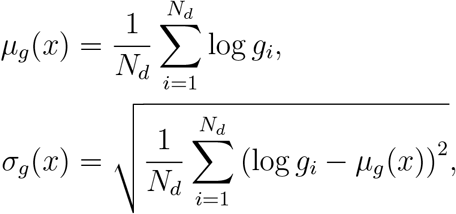

where 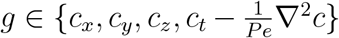 and *N*_*d*_ denotes the number of data points at a given time. Accordingly, we discard data points where any of the gradients lie outside the range *µ*_*g*_ ± 3*σ*_*g*_.

Additionally, analysis of the synthetic data (see Figure B.10(C)) shows that concentration values below 10^−2^ tend to diverge from the target. To avoid introducing unreliable supervision, we also discard concentration values below 10^−1^ when learning the velocity field. We emphasize that these low-concentration regions provide minimal information for velocity reconstruction, and their exclusion prevents the propagation of weak or ambiguous signals into the PDE-constrained learning stage. Notably, this filtering step is applied exclusively during velocity inference; the concentration model is trained using unfiltered concentration data.

##### Training

**Step 2** We begin the training stage by refining the initial pressure field. During this step, we fix the concentration and permeability parameters ***θ***_*c*_ and ***θ***_*K*_ learned in the initialization stage, and update only the pressure parameters ***θ***_*P*_. Pressure refinement is performed by enforcing the physical laws, aiming to minimize the advection-diffusion and conservation of mass residuals through the equation loss:

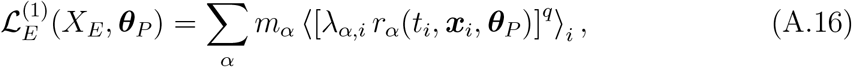

where *α* = {AD, CM} indexes the advection-diffusion and conservation of mass equation residuals, respectively. As in the previous cases, *m*_*α*_ is the global weight balancing the contribution of each term, and *λ*_*α,i*_ is the corresponding residual-based attention (RBA) weight. In this step, we use *q* = 2, inducing a squared norm appropriate for the initial training stage. Our analysis of the concentration revealed that the concentration and its gradients have a fast decay after the first few minutes of an experiment (see Figure A.8(A)). Therefore, we further split the time domain into three intervals and apply three separate losses to aid the model to learn the information from the low concentration gradients. We train the pressure network for *N*_*s*2_ = 50000 ADAMw [64] iterations.

**Step 3** After reducing the advection-diffusion residuals through pressure refinement, we proceed to jointly refine the permeability field. At this stage the pressure and permeability parameters ***θ***_*K*_, and ***θ***_*P*_ are updated following equation A.16. We train the pressure and permeability networks for *N*_*s*3_ = 150000 ADAMw [64] iterations.

**Step 4** At this stage, all network parameters, including ***θ***_*c*_, ***θ***_*K*_, and ***θ***_*P*_, are updated by minimizing a combined loss that balances both the equation residuals and the mismatch with the experimental concentration data. The data loss *ℒ*_*D*_, which constrains only the concentration field, is defined as:

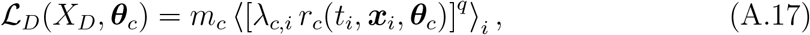

where 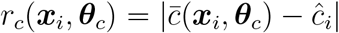 is the residual between the reconstructed mean concentration and the experimental observation *ĉ*_*i*_ at location ***x***_*i*_, *m*_*c*_ is the global weight for the concentration term, *λ*_*c,i*_ is the corresponding local weight, and *q* = 2 is the exponent inducing a quadratic loss. The combined loss for this full training stage is given by:

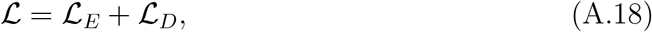

where *ℒ*_*D*_ is the data loss penalizing the mismatch with the experimental concentration data (see equation A.17) and *ℒ*_*E*_ is the equation loss enforcing the advection-diffusion and conservation of mass constraints:

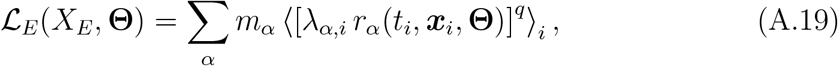

Notice that the main difference with equation A.16 is that we are now optimizing all the parameters **Θ** = {***θ***_*P*_, ***θ***_*c*_, ***θ***_*K*_}.

We train the pressure, permeability, and concentration networks for *N*_*s*4_ = 150000 ADAMw [64] iterations.

**Step 5** Finally, since a quadratic loss (*q* = 2) tends to over-smooth the loss landscape, especially in the later stages of training, we include a refinement stage that uses *q* = 1. This choice promotes sharper gradients and improves the model’s ability to capture fine-scale details. This strategy was introduced in [37], where it was experimentally shown to improve the performance of PIML models. We train the pressure, permeability, and concentration networks for *N*_*s*5_ = 50000 ADAMw [64] iterations.

Notice that the final stages correct the initial field estimates and ensure that the model produces physically consistent solutions that also match the observed concentration data.

### Appendix A.3. Residual-Based Attention Methods

#### Appendix A.3.1. Residual-Based Attention weights (RBA)

One of the main challenges in training neural networks is that residuals (i.e., pointwise errors) may be overlooked when computing a cumulative loss function [42, 65]. To address this issue, we employ residual-based attention (RBA) [43] as local weights (*λ*_*α,i*_), which help balance the point-wise contribution of each residual term *α*. RBA acts as an attention mechanism, guiding the optimizer to focus on spatiotemporal regions where the residuals remain high [42, 43]. The update rule for an RBA weight *λ*_*α,i*_, associated with loss term *α* and point *x*_*i*_, is based on the exponentially weighted moving average of the residuals:

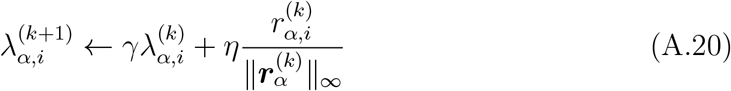

where 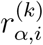 is the residual for loss term *α* at point *i*, 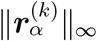 is the maximum residual at iteration *k, η* is the learning rate for the weights, and *γ* is a memory term.

#### Appendix A.3.2. Residual-Based Attention with Resampling (RBA-R)

since RBA weights are updated based on the residuals computed at each specific iteration, inconsistencies can arise for large data sets trained in batches. To mitigate this issue, we leverage the RBA weights to resample critical points directly [37]. Because the weights contain historical information about regions with persistent high error, they can be used to construct a stable sampling probability density function (PDF) [66]:

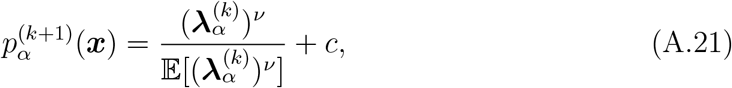

where 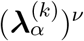 is the vector of RBA weights raised to a power *ν* that controls the sharpness of the distribution, and *c >* 0 is a small scalar ensuring that all points retain a nonzero sampling probability. This combined approach is referred to as RBA with resampling (RBA-R).

#### Appendix A.3.3. Time-dependent RBA (TD-RBA)

In our framework, we apply RBA-R to all loss terms. However, the advection-diffusion equation presents an additional critical challenge: as shown in Figure A.8, the concentration gradients vary by several orders of magnitude over time. This variability means that time points with large gradients can disproportionately dominate the optimization process, preventing the model from learning from the weaker signals present at other times.

To address this imbalance, we introduced our novel optimization method, Time-dependent RBA (TD-RBA). The core idea is to introduce a time-dependent scaling factor, *C*(*t*), to ensure that the residuals at all times contribute equally during training. The modified loss for the advection-diffusion equation takes the form:

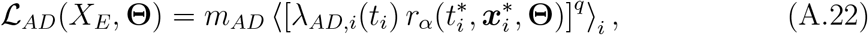

where the TD-RBA weight is 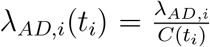. The scaling factor *C*(*t*_i_) is defined by the maximum magnitude of the advection-diffusion equation’s components across the spatial domain Ω_*x*_ at time 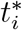:

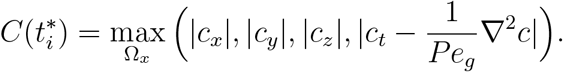

Since this scaling factor is computed from the clean concentration field (which is learned in an early training stage), it is calculated only once. As shown in Figure A.8B, this rescaling successfully maintains the maximum gradient values within a stable range across all times. Our ablation study (Table B.2) demonstrates that this physics-guided normalization is critical for reducing modeling errors.

**Figure A.8:**
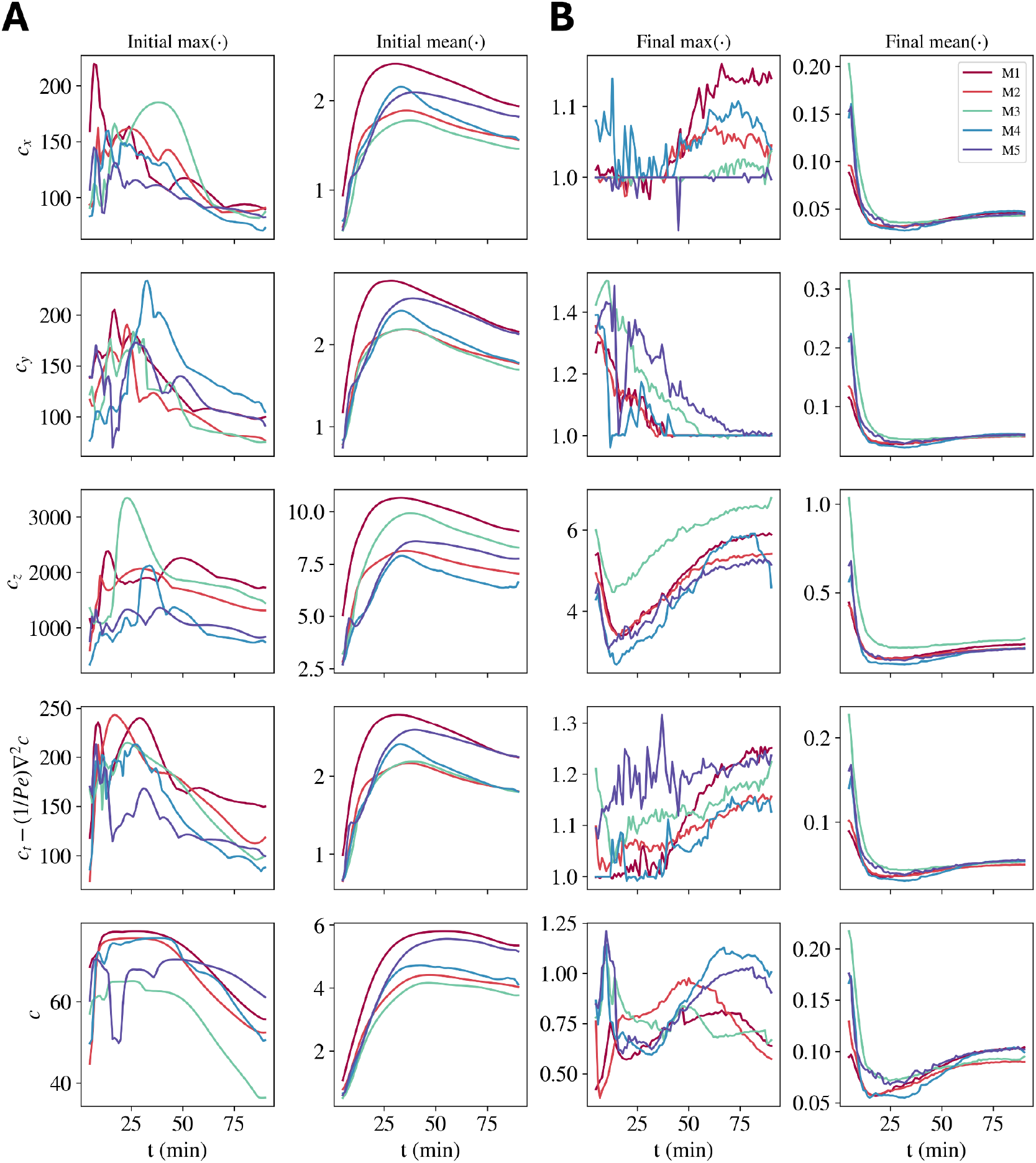
Effect of filtering and time-dependent residual-based attention (TD-RBA) scaling on the concentration gradients. (A) Time evolution of the maximum and spatially-averaged concentration gradients before filtering and scaling: *c*_*x*_, *c*_*y*_, *c*_*z*_, 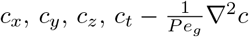, and *c*. Results are shown for five different mice (M1–M5). Notice the large fluctuations and substantial variability across times and mice, spanning several orders of magnitude. (B) Time evolution of the scaled gradients after filtering and applying TD-RBA scaling. The maximum values are of order one, and means are consistent across all mice. This rescaling ensures uniform contribution of the residuals across times, improving the stability of the velocity inference process.

**Figure B.9:**
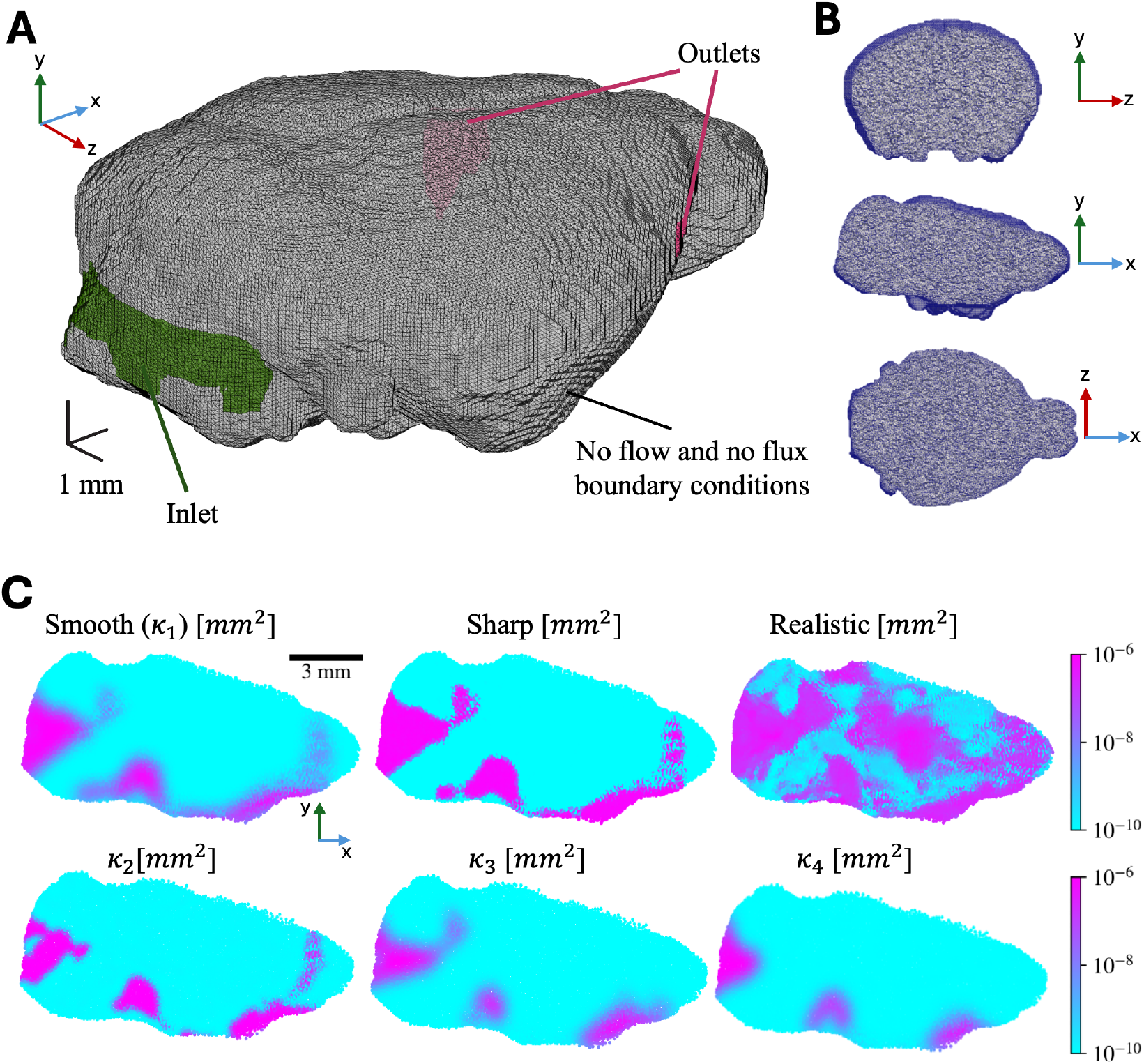
Tetrahedral mesh and permeability maps for producing synthetic data. **(A)** Finite element mesh used for simulations, consisting of approximately 2 million tetrahedral elements. The inlet, outlets, and no-flow boundary conditions are indicated. **(B)** Cross-sectional views of the mesh. **(C)** Permeability maps used in simulations: “Smooth” with gradual transitions, “Sharp” with abrupt interfaces, and “realistic” with a complex map that combines sharp and irregular transitions. Maps *κ*_1_ − *κ*_4_ were used as different initial guesses, allowing uncertainty quantification.

## Appendix B. Numerical Simulations for Synthetic Data

To generate the synthetic data, we used the finite element method code COMSOL Multiphysics 6.2 to perform the numerical simulations. The simulations were performed in two steps: the first step was solving for the fluid velocity field, and the second step was calculating the concentration by solving the advection-diffusion equation. The brain was considered to be a porous medium in the simulations, and the fluid motion in the domain was governed by Darcy’s law and continuity; see equations A.3 and A.2, respectively. In all simulations, we used *µ* = 0.7 × 10^−3^ Pa · s, the viscosity of water at 37^°^C.

The basic equation governing the concentration *C*(**x**, *t*) of a passive solute in the brain is the *advection-diffusion equation* (see equation A.1). We first validated our numerical methods by comparing the numerical results with analytical solutions of one-dimensional advection-diffusion problems, obtaining good agreement. Mesh sensitivity studies were also performed to ensure that the meshes were sufficiently fine to resolve the computational domains and that the numerical results did not change substantially when the mesh size was decreased further. In the simulation results shown in this study, the total number of tetrahedral elements is approximately 2 million, and the mesh size ranges from 0.05 mm to 0.2 mm. The quintic finite element was used when solving the Darcy equation, and the quadratic finite element was used when solving the advection-diffusion equation.

The 3D domain boundaries were obtained from the segmentation of the brain region from Mouse 1 (see Figure B.9(A)). In the simulations, we defined an inlet region close to and approximately the same size as the cisterna magna, and we placed two outlets on the domain surface at locations where tracer leaves the segmented brain region in the experimental data. When solving Darcy’s law for the fluid velocity field, a constant pressure of 7.5 kPa was applied at the inlet, and zero pressure was applied at the outlets. When solving the advection-diffusion equation, a constant tracer concentration was applied at the inlet for the first 5 minutes, then set to 0 for the remainder of the simulation. All other domain boundaries are treated as no-flow (i.e., **n** · **u** = 0, where **n** is the surface normal) when solving Darcy’s law, and as no-flux (i.e., **n** · ∇*C* = 0) when solving the advection-diffusion equation (see Figure B.9(A)).

To simulate DCE-MRI, we interpolated the concentration field onto a 0.1 mm square grid (the same resolution as the DCE-MRI data) at 1-minute increments for 90 minutes.

### Appendix B.1. Concentration Reconstruction

We train our models using 50% of the available concentration data, which is used both to learn the concentration field and to extract aleatoric uncertainty via the denoising module composed of *NN*_*c*_ and *NN*_*σ*_. Model performance is then evaluated on the remaining unseen data. As shown in Table B.1, the validation error varies depending on the underlying permeability map.

**Figure B.10:**
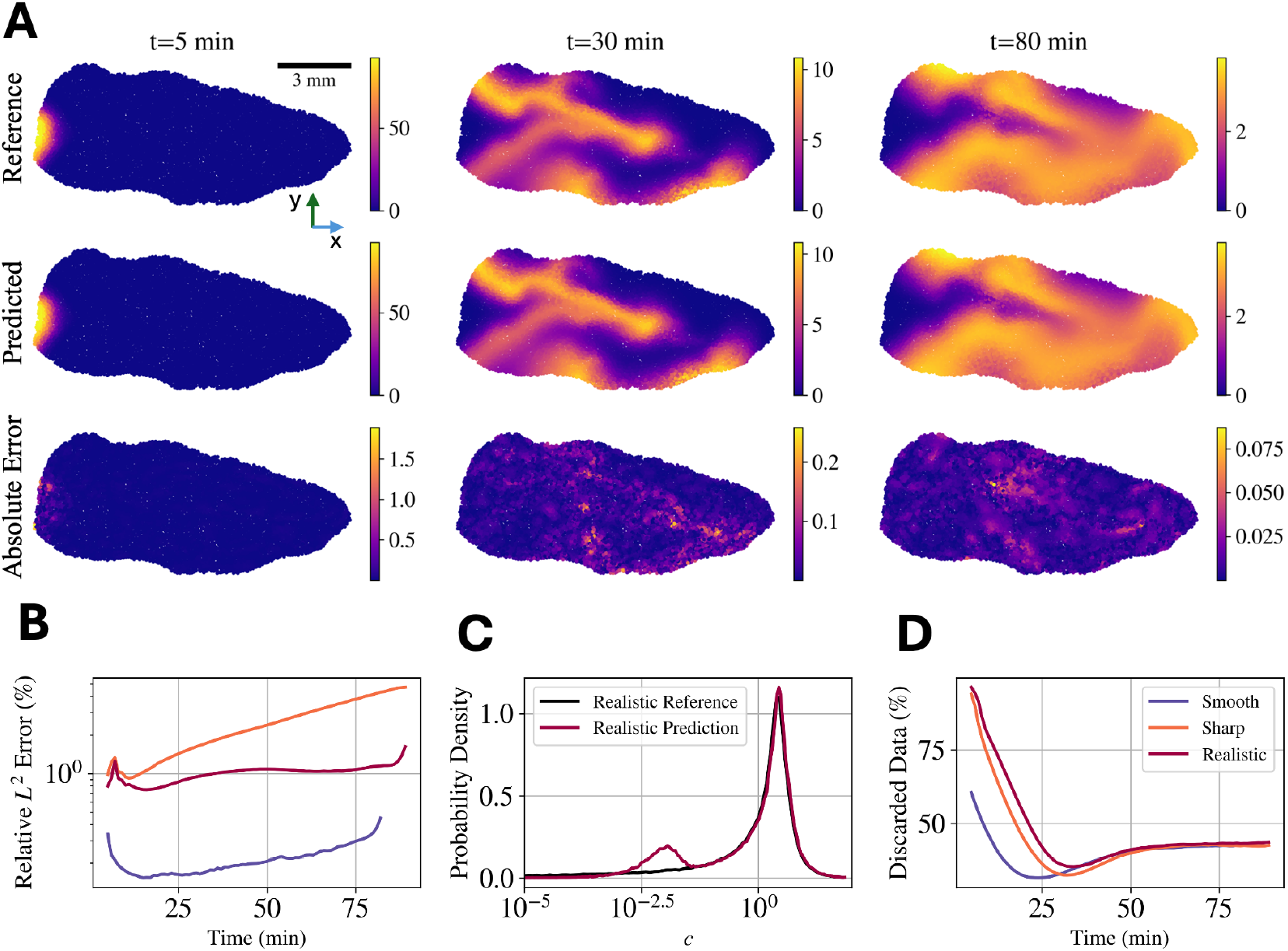
Reconstruction of concentration using synthetic data. (A) Tracer concentration in the mid-sagittal plane for the “realistic” synthetic case, at times *t* = 5, 30, and 80 minutes, as measured and as inferred. The reconstructed concentration shows spatially coherent transport. The absolute error, also shown, is greatest in regions of high concentration, particularly at early times, and decreases as the tracer spreads. (B) Relative *L*^2^ error between inferred and measured concentrations varying over time for synthetic data with “smooth”, “sharp”, and “realistic” permeability. In the “realistic” case, the error usually remains below 2%, with an overall average near 1%. (C) Probability density functions of inferred and measured concentrations for the “realistic” case. The model accurately captures the full distribution across six orders of magnitude, though a mismatch appears in the low-concentration regime (*c <* 10^−3^) due to weak signals. (D) Percentage of data discarded, based on thresholds tied to sensitivity, varying over time, for all synthetic cases. The discard rate is highest (up to 90%) for the “realistic” case and at early times; it decreases to about 40% as the tracer spreads.

**Table B.1:** Reconstruction performance on synthetic data. Relative *L*^2^ errors and Wasserstein distances (*W*_*d*_) for different synthetic permeability maps. Smooth, sharp, and realistic cases evaluate the model’s performance under increasing structural complexity. While *L*^2^ errors capture pointwise accuracy, *W*_*d*_ highlights discrepancies in the recovered velocity distributions.

| Permeability map | Relative $L^2$ Error (%) | | | | $W_d \cdot 10^2$ | | |
| --- | --- | --- | --- | --- | --- | --- | --- |
| | $c$ | $\ \mathbf{u}\ $ | $P$ | $\kappa$ | $\log_{10}\vert u\vert $ | $\log_{10}\vert v\vert $ | $\log_{10}\vert w\vert $ |
| Smooth | 0.9 | 12.7 | 76.1 | 89.4 | 4.0 | 5.4 | 6.3 |
| Sharp | 2.3 | 26.2 | 77.1 | 63.8 | 14.1 | 10.9 | 13.7 |
| Realistic | 1.0 | 36.0 | 41.9 | 59.1 | 18.0 | 15.7 | 15.5 |

For the “smooth” map, the model achieves a relative *L*^2^ error of 0.2%; for the “sharp” map, performance decreases to 2%; and for the “realistic” map, the error is approximately 1%. Figure B.10(A) shows the reference, prediction, and absolute error of the concentration at three representative times.

Additionally, Figure B.10(B) shows that the error remains consistent across all times, with a slight degradation in performance at the beginning and end of the time domain. Figure B.10(C) presents the probability density function (PDF) of the concentration, illustrating that, even though the concentration in the “realistic” data set spans over six orders of magnitude, the model successfully captures the distribution.

Due to the wide dynamic range of the concentration field, we observe a mismatch between reconstructed and reference values in the low-concentration regime (*c <* 10^−4^), as shown in Figure B.10(C). Despite this, the concentration model is trained using all available observations, including those in the low-concentration regime. However, when inferring the velocity field, we introduce a post-processing step to discard regions with low concentration and weak gradients based on the denoised data. This filtering step is intended to prevent the propagation of low-information content into the PDE-constrained velocity learning stage. Specifically, we define a threshold on both the concentration and its gradient to identify regions that provide an insufficient signal for supervising the velocity field. These regions are explicitly excluded from the velocity inference process, as their inclusion could degrade performance by introducing weak or ambiguous supervision. The specific details of the filtering process are described in Appendix A.2.

We observe that, most of the time, at least 40% of the domain falls below this threshold (see Figure B.10(D)). Interestingly, the percentage of discarded data increases to as much as 90% during early time points, aligning with regions where the model has the greatest difficulty reconstructing the concentration field (see Figure B.10(B)).

### Appendix B.2. Inferring hidden 3D fields

#### Appendix B.2.1. 3D velocity Inference

Since the true permeability is unknown in real data sets, we evaluate the robustness of velocity inference in the synthetic validation by using mismatched permeability estimates. Specifically, for the “smooth” case, we use the “sharp” permeability map as the initial estimate. Conversely, for both the “sharp” and “realistic” cases, we use the “smooth” map as the initial guess.

**Figure B.11:**
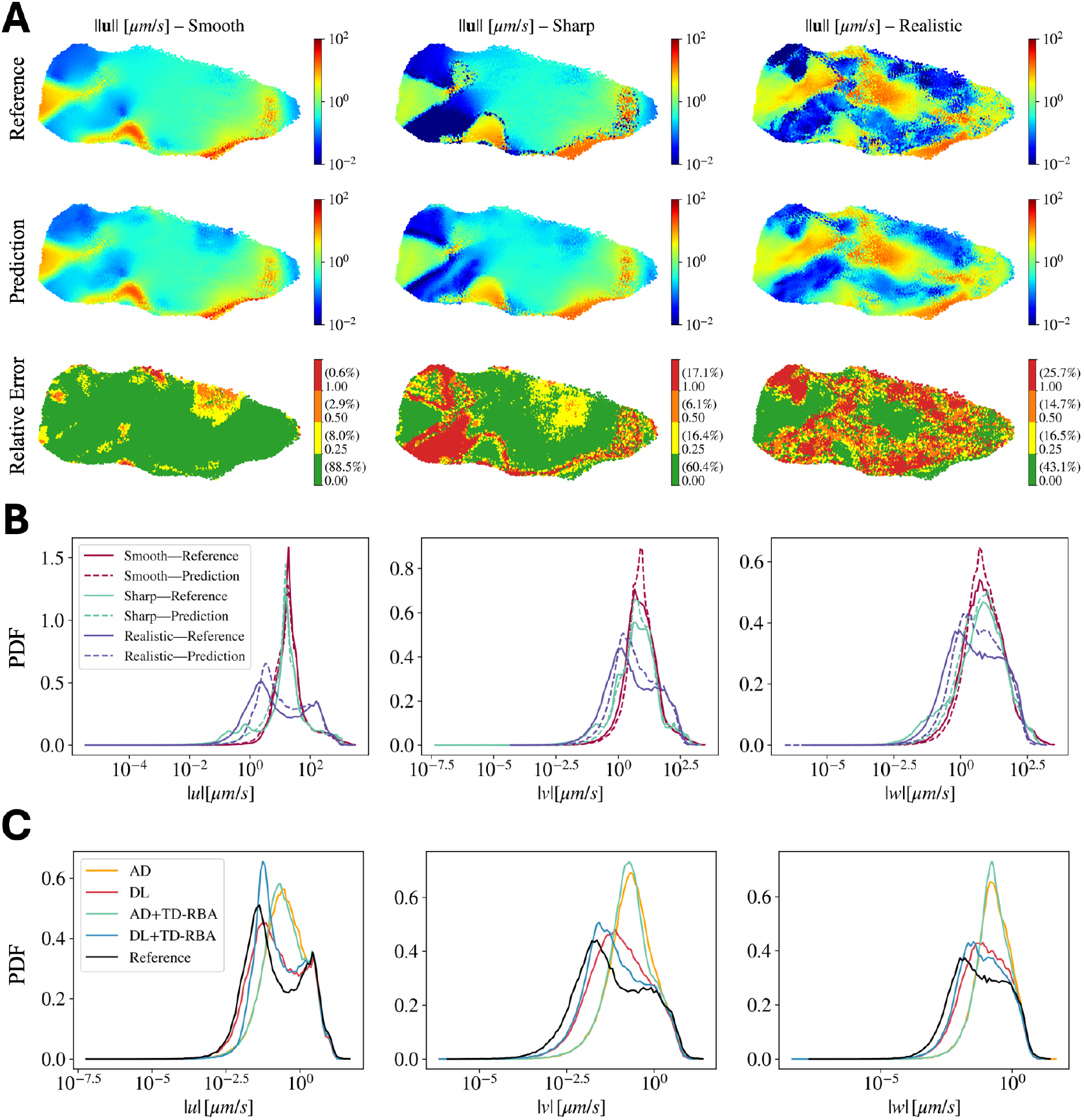
Velocity inference using synthetic data. (A) Reference, prediction, and relative error of the velocity magnitude ∥**u**∥ for three synthetic velocity fields: “smooth”, “sharp”, and “realistic”. While accuracy is high for the “smooth” case, errors increase for the more complex fields, concentrating in low-velocity regions and at transition zones. (B) Probability density functions (PDFs) of velocity component magnitudes. The predicted distributions (dashed) accurately capture the reference (solid), including the bimodal structure induced by the “realistic” permeability, across five orders of magnitude. (C) PDFs from an ablation study on the “realistic” dataset. Only models enforcing Darcy’s law (DL) recover the bimodal velocity distributions. Models using only the advection-diffusion equation (AD) predict a unimodal, averaged solution, likely due to convergence to poor local minima.

**Figure B.12:**
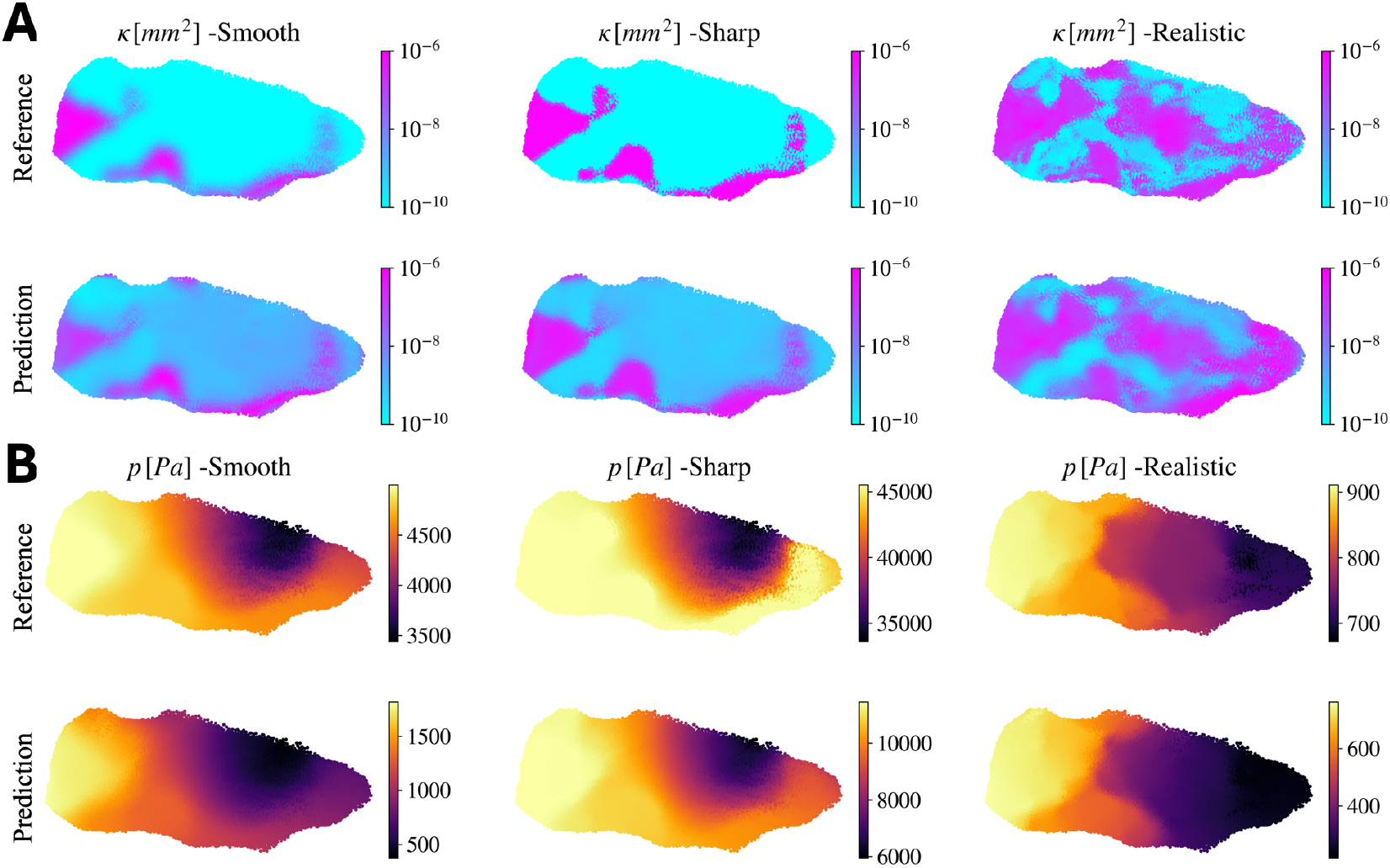
Pressure and permeability estimation using synthetic data. (A) Reference and predicted permeability fields for synthetic data with “smooth”, “sharp”, and “realistic” permeabilities. Models successfully capture the spatial structure and magnitude of the permeability in all cases. (B) Reference and predicted pressure fields for the same data. Models also accurately reproduce the pressure map.

Figure B.11(A) shows the reference, predictions, and point-wise relative error for the three synthetic cases. In all scenarios, the velocity magnitude spans more than four orders of magnitude. Notably, in the “smooth” case, the model achieves less than 25% relative error in nearly 90% of the domain. As shown in Table B.1, the relative *L*^2^ error for the velocity magnitude ∥**u**∥ is 12.7%.

The “sharp” permeability map, by contrast, induces abrupt transitions in the velocity field and regions where the velocity becomes extremely small. Despite these challenges, the model still achieves less than 25% relative error in approximately 60% of the domain, with relative *L*^2^ error for the velocity magnitude of 26.2%. As shown in Figure B.11(A), the error is primarily concentrated in regions with very low velocity (∥**u**∥ ≤ 10^−2^µm/s) and at transitions between high and low velocities.

This pattern persists in the “realistic” case, which contains a larger fraction of low-velocity regions. For this case, the relative *L*^2^ errors for the velocity magnitude reach36%. However, as shown in Figure B.11(A), the errors are primarily concentrated in low-velocity regions. Nevertheless, about 40% of the domain still exhibits relative errors less than 25%. This highlights an important limitation of our method: it struggles to reconstruct extremely low velocities. This behavior is expected, as very slow velocities result in minimal tracer transport (see Figure B.10(C)), which limits the information available for accurate reconstruction.

Finally, Figure B.11(B) shows that, in the “sharp” and “realistic” cases, the velocity components span a range from 10^−2^ µm/s to 10^2^ µm/s, with most values concentrated between 10^−2.5^ µm/s and 10^−1.5^ µm/s. Despite this challenging distribution, the predicted velocity histograms closely match the reference distributions, demonstrating the model’s ability to recover the full range of values. Moreover, in the “realistic” case, the complexity of the permeability map induces bimodal distributions in the velocity components, which are also successfully captured by the model.

#### Appendix B.2.2. Permeability and Pressure Estimates

As shown in Figure B.12, models successfully capture the overall spatial structure of both pressure and permeability across all three synthetic cases. The predicted fields exhibit strong agreement with the reference fields, preserving key structural features and regional contrasts. However, while the spatial trends are well recovered, the absolute magnitudes of the predicted fields deviate from the ground truth. This discrepancy is reflected in the relative *L*^2^ errors reported in Table B.1, where pressure errors range from 76.1% to 48.6%, and permeability errors from 89.4% to 51.3% across the “smooth”, “sharp”, and “realistic” cases.

These high errors in magnitude stem from the inherent ambiguity in estimating pressure and permeability from velocity-constrained data. According to Darcy’s law, **u** = −*K*∇*P*, the same velocity field can result from many different combinations of *K* and ∇*P*. Since the concentration data only constrains the velocity field through the advection–diffusion equation, it cannot uniquely determine either *K* or *P*. The improved accuracy observed in the “realistic” case likely arises from the added flow complexity, which imposes stronger implicit constraints and reduces the space of admissible solutions. As discussed in Appendix B, some permeability initial guesses induce considerably lower errors; nonetheless, the inverse problem remains ill-posed, and the inferred fields should be interpreted as plausible and physically consistent, but not unique.

#### Appendix B.3. Ablation Study

The proposed framework incorporates several key enhancements, which can be decomposed into the following components: (1) a specialized architecture that strictly enforces Darcy’s law (DL), and (2) a time-dependent residual-based attention (TS-RBA) weighting scheme. To evaluate the contribution of each component, we performed an ablation study comparing the DL-based architecture against a baseline architecture (AD), which consists of a single multi-layer perceptron (MLP) that predicts the velocity components (*u, v, w*) without explicitly enforcing Darcy’s law. All models were trained using the same sequential strategy and denoising module to ensure a fair comparison. Table B.2 summarizes the results of the ablation study across different model configurations. Notice that the DL-based formulation is crucial to reduce the errors, especially in the w-component, which is harder to recover due to its considerably smaller magnitude (See Figure B.11). Moreover, the Wasserstein distance (*W*_*d*_) reveals a critical limitation of the AD architecture: despite similar average errors, it fails to capture the correct velocity distributions, as reflected by substantially higher *W*_*d*_ values across all components. This limitation is further illustrated in Figure B.11(A), which shows the distributions of the velocity component magnitudes |*u*|, |*v*|, and |*w*|. Only the DL-based models recover the characteristic bimodal structure of the velocity distribution, while the AD models tend to collapse toward intermediate values between the two peaks, effectively predicting an averaged solution, which may reflect convergence to a poor local minimum.

**Table B.2:** Ablation study of model architecture and training strategies. The baseline architecture (AD) uses a direct MLP to predict velocity without enforcing Darcy’s law, while the DL architecture encodes this constraint explicitly. All models share the same denoising module and sequential training for fair comparison. Relative *L*^2^ errors measure average accuracy, and Wasserstein distances (*W*_*d*_) assess the quality of the reconstructed velocity distributions. While AD achieves similar relative *L*^2^ errors, it fails to capture the correct distributions. DL reduces *W*_*d*_ substantially, and TD-RBA further improves the recovery of velocity, pressure, and permeability.

| Architecture | Relative $L^2$ Error (%) | | | | | | $W_d \cdot 10^{-2}$ | | |
| --- | --- | --- | --- | --- | --- | --- | --- | --- | --- |
| | $c$ | $u$ | $v$ | $w$ | $P$ | $\kappa$ | $\log_{10}\vert u\vert $ | $\log_{10}\vert v\vert $ | $\log_{10}\vert w\vert $ |
| AD | 0.9 | 35.0 | 45.8 | 71.1 | - | - | 34.5 | 54.4 | 52.8 |
| DL | 0.9 | 43.2 | 56.5 | 60.5 | 80.0 | 85.8 | 9.2 | 23.1 | 21.2 |
| AD+Seq+TD-RBA | 1.0 | 36.9 | 45.9 | 54.1 | - | - | 33.2 | 51.5 | 51.4 |
| DL+Seq+TD-RBA | 1.0 | 35.5 | 44.2 | 48.0 | 43.9 | 51.3 | 18.0 | 15.7 | 15.5 |

The DL architecture consistently improves the distributional accuracy, substantially reducing the Wasserstein distance, especially for the dominant velocity component. Furthermore, the inclusion of the TD-RBA weighting scheme provides additional gains, particularly when combined with sequential training. Notably, TD-RBA not only improves velocity reconstruction but also facilitates the recovery of pressure and permeability fields, which are otherwise unstable without these enhancements.

## Appendix C. Real Data Additional Results

### Appendix C.1. Permeability and Pressure Estimation

To enable consistent visualization across mice, we apply a normalization step that enforces a common maximum permeability of 10^−6^ mm^2^ and displaces each pressure field so that all models have the same mean. This transformation is justified because

**Figure C.13:**
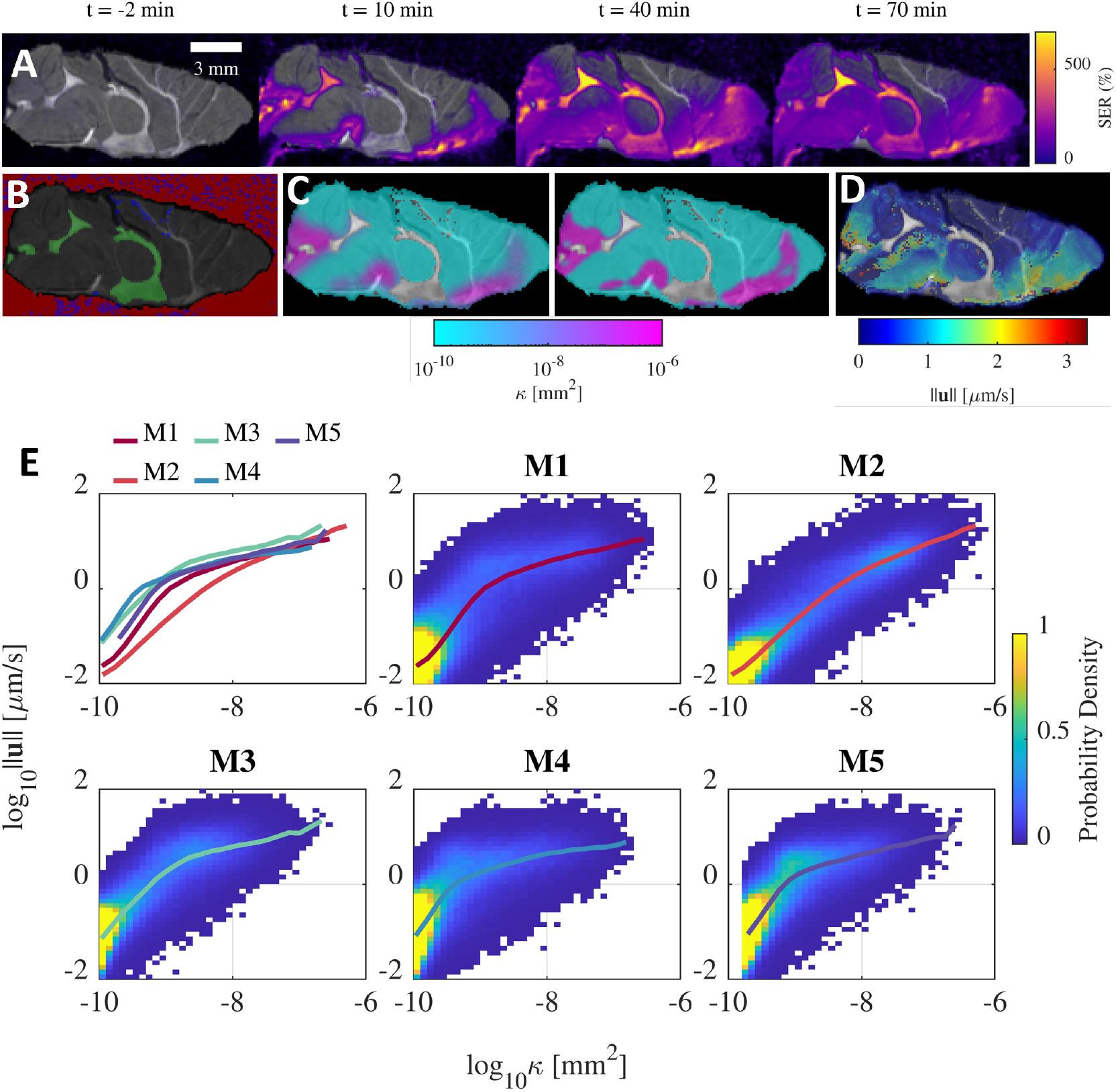
DCE-MRI data and preprocessing. (A) Signal enhancement ratio (SER) over time for Mouse 1 after gadobutrol injection. (B) Excluded regions for analysis. (C) Initial permeability estimates (“smooth” and “sharp”) derived from early tracer concentration. (D) Initial velocity estimate from front tracking. (E) **Velocity magnitude and permeability are closely coupled**. The bivariate probability density (heat map) and conditional average (solid line) for each mouse confirm that high velocities occur in regions with high inferred permeability.

**Table C.3:** Median velocity in deep brain regions. Median velocity magnitude ||**u**|| [µm/s] for selected deep brain regions and the sagittal sinus. Data is shown for five mice (M1-M5) along with the median and standard deviation across all mice.

|  | Hippocampus | Caudate | Thalamus | Midbrain | Sagittal Sinus |
| --- | --- | --- | --- | --- | --- |
| M1 | 0.043 | 0.029 | 0.028 | 0.042 | 0.097 |
| M2 | 0.025 | 0.030 | 0.024 | 0.035 | 0.515 |
| M3 | 0.104 | 0.066 | 0.074 | 0.115 | 1.050 |
| M4 | 0.154 | 0.097 | 0.060 | 0.103 | 1.039 |
| M5 | 0.205 | 0.123 | 0.110 | 0.128 | 0.578 |
| Median | 0.106 | 0.069 | 0.059 | 0.085 | 0.656 |
| Deviation | 0.067 | 0.037 | 0.032 | 0.039 | 0.358 |

**Table C.4:**
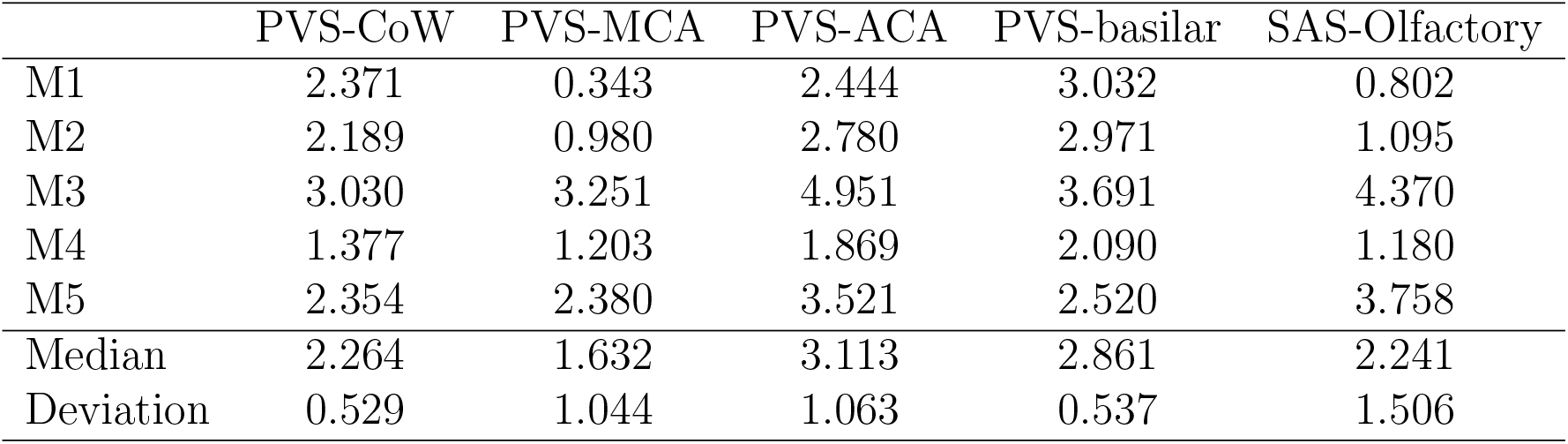
Median velocity in perivascular and subarachnoid spaces. Median velocity magnitude ||**u**|| [µm/s] for selected perivascular spaces (PVS) and the subarachnoid space (SAS) of the olfactory bulb. Acronyms: CoW, Circle of Willis; MCA, middle cerebral artery; ACA, anterior cerebral artery. Data is shown for five mice (M1-M5) along with the median and standard deviation across all mice.

|  | PVS-CoW | PVS-MCA | PVS-ACA | PVS-basilar | SAS-Olfactory |
| --- | --- | --- | --- | --- | --- |
| M1 | 2.371 | 0.343 | 2.444 | 3.032 | 0.802 |
| M2 | 2.189 | 0.980 | 2.780 | 2.971 | 1.095 |
| M3 | 3.030 | 3.251 | 4.951 | 3.691 | 4.370 |
| M4 | 1.377 | 1.203 | 1.869 | 2.090 | 1.180 |
| M5 | 2.354 | 2.380 | 3.521 | 2.520 | 3.758 |
| Median | 2.264 | 1.632 | 3.113 | 2.861 | 2.241 |
| Deviation | 0.529 | 1.044 | 1.063 | 0.537 | 1.506 |

Darcy’s law, ***u*** = −*K*∇*P*, is invariant under the rescaling and translation 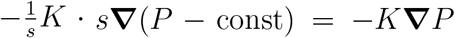. While this normalization allows for meaningful visual comparison, it does not resolve the fundamental non-uniqueness in recovering *K* and *P* from velocity alone. Consequently, the inferred fields should be interpreted as plausible and physically consistent estimates, not as unique reconstructions.

**Table C.5:** Median permeability in deep brain regions. Median permeability *κ* (× 10^−10^ [mm^2^]) for selected deep brain regions and the sagittal sinus. Data is shown for five mice (M1-M5) along with the median and standard deviation across all mice.

|  | Hippocampus | Caudate | Thalamus | Midbrain | Sagittal Sinus |
| --- | --- | --- | --- | --- | --- |
| M1 | 3.86 | 3.73 | 3.95 | 4.35 | 5.37 |
| M2 | 2.73 | 2.45 | 2.54 | 3.38 | 17.8 |
| M3 | 4.35 | 3.93 | 3.96 | 4.25 | 16.4 |
| M4 | 6.46 | 6.11 | 5.96 | 6.78 | 10.8 |
| M5 | 7.81 | 6.82 | 6.50 | 6.87 | 13.0 |
| Median | 4.35 | 3.93 | 3.96 | 4.35 | 13.0 |
| Deviation | 1.84 | 1.62 | 1.45 | 1.43 | 4.42 |

**Table C.6:** Median permeability in perivascular and subarachnoid spaces. Median permeability *κ* (×10^−10^ [mm^2^]) for selected perivascular spaces (PVS) and the subarachnoid space (SAS) of the olfactory bulb. Acronyms: CoW, Circle of Willis; MCA, middle cerebral artery; ACA, anterior cerebral artery. Data is shown for five mice (M1-M5) along with the median and standard deviation across all mice.

|  | PVS-CoW | PVS-MCA | PVS-ACA | PVS-basilar | SAS-Olfactory |
| --- | --- | --- | --- | --- | --- |
| M1 | 110 | 11.5 | 155 | 31.9 | 41.8 |
| M2 | 148 | 45.2 | 281 | 78.1 | 79.9 |
| M3 | 49.3 | 44.3 | 135 | 26.4 | 123 |
| M4 | 32.4 | 27.6 | 127 | 35.1 | 22.6 |
| M5 | 55.4 | 69.0 | 153 | 23.5 | 169 |
| Median | 55.4 | 44.3 | 153 | 31.9 | 79.9 |
| Deviation | 43.2 | 19.2 | 56.3 | 20.0 | 53.2 |

### Appendix C.2. Uncertainty Quantification

Psaros et al. [67] identified three main sources of uncertainty in PIML: aleatoric uncertainty, which originates from noise or sparsity in the data; epistemic uncertainty, which stems from limitations in the model architecture, initialization, or assumptions; and model uncertainty, which arises from assumptions in the governing equations. In our approach, aleatoric uncertainty in the concentration field is estimated directly from the experimental data using a denoising network trained with a negative log-likelihood loss, providing uncertainty estimates for concentration measurements (Figure 2(A)).

Epistemic uncertainty, which reflects the sensitivity of model predictions to architectural choices and parameter initialization, is traditionally assessed by training multiple models with different random initializations[68]. In our framework, however, due to the use of a pre-training stage, the initial permeability guess effectively defines the parameter initialization for the velocity inference. To quantify this uncertainty, we adopt an ensemble-of-models (EoM) approach [10, 41, 69], where multiple models are trained independently, each initialized with a different permeability guess. Following [41], we approximate the ensemble prediction as a Gaussian distribution, with the mean and variance given by the empirical mean and variance of the individual model outputs, as defined in equations C.1 and C.2, respectively.

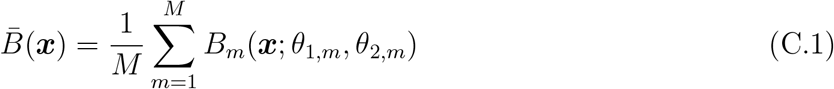

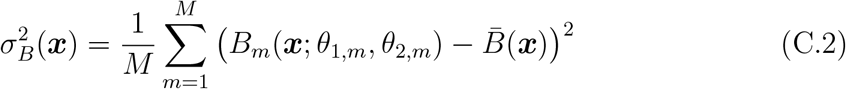

where *B u, v, w, P, K* denotes the physical quantities of interest: the velocity components (*u, v, w*), pressure (*P*), and permeability (*K*).

We apply this method to both synthetic (“realistic” map) and real data sets (M1) using four different initial permeability maps (see Figure B.9). One of our key assumptions is that we can derive a reasonable initial guess of the permeability map from the early tracer distribution. To evaluate the sensitivity of the inferred results on the initial guess, we compare results using *κ*_3_ and *κ*_4_ permeability fields derived from Mice 3 and 4, respectively, which differ substantially from the standard guess *κ*_1_ (from Mouse 1). Additionally, to isolate the influence of structural sharpness, we include the map *κ*_2_, which is derived from the Mouse 1 data and contains abrupt transitions designed to challenge the learning process. This ensemble analysis enables a systematic evaluation of the model’s sensitivity to permeability initialization and the robustness of the inferred fields.

Finally, to provide a coarse estimate of the modelling uncertainty, we include results from a baseline model that infers the velocity field using only the advection-diffusion equation. This model is not physically well-posed, as the combination of advection-diffusion and mass conservation involves four coupled unknowns, but it serves to illustrate the limitations of relying solely on tracer transport without incorporating additional physical constraints.

**Figure C.14:**
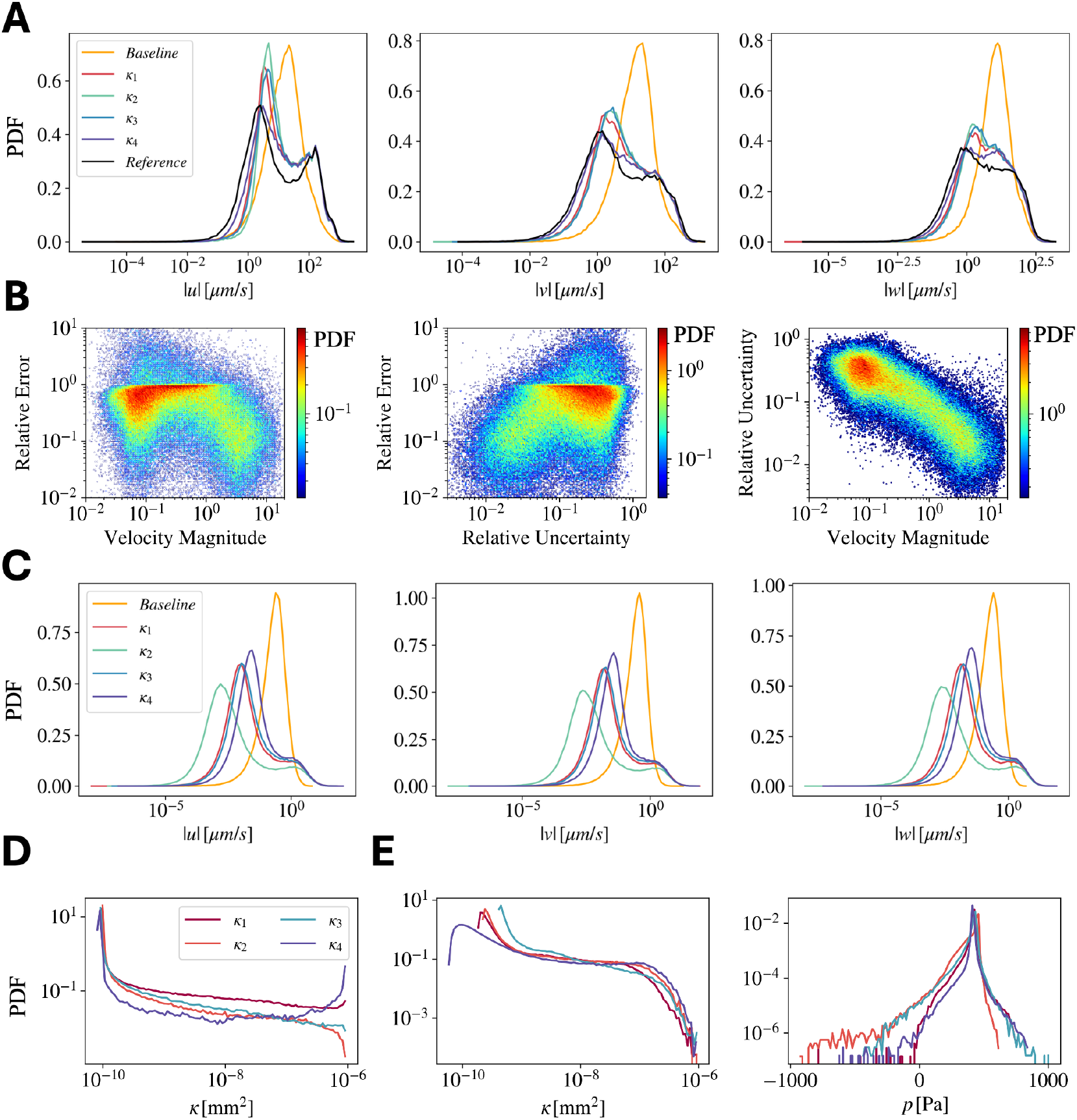
Distributions and uncertainty quantification. (A) Probability density functions (PDFs) of velocity component magnitudes for the “realistic” synthetic data. Models enforcing Darcy’s law successfully capture the bimodal structure of the reference data, while the baseline model (which only enforces advection-diffusion and continuity) fails. (B) Joint distributions for the synthetic data show that relative error is highest in slow regions and that predicted uncertainty strongly correlates with error. (C) PDFs of velocity component magnitudes for Mouse 1. (D) PDFs of the initial permeability guesses. (E) PDFs of the inferred permeability and pressure for Mouse 1, using different initial permeability guesses.

**Table C.7:**
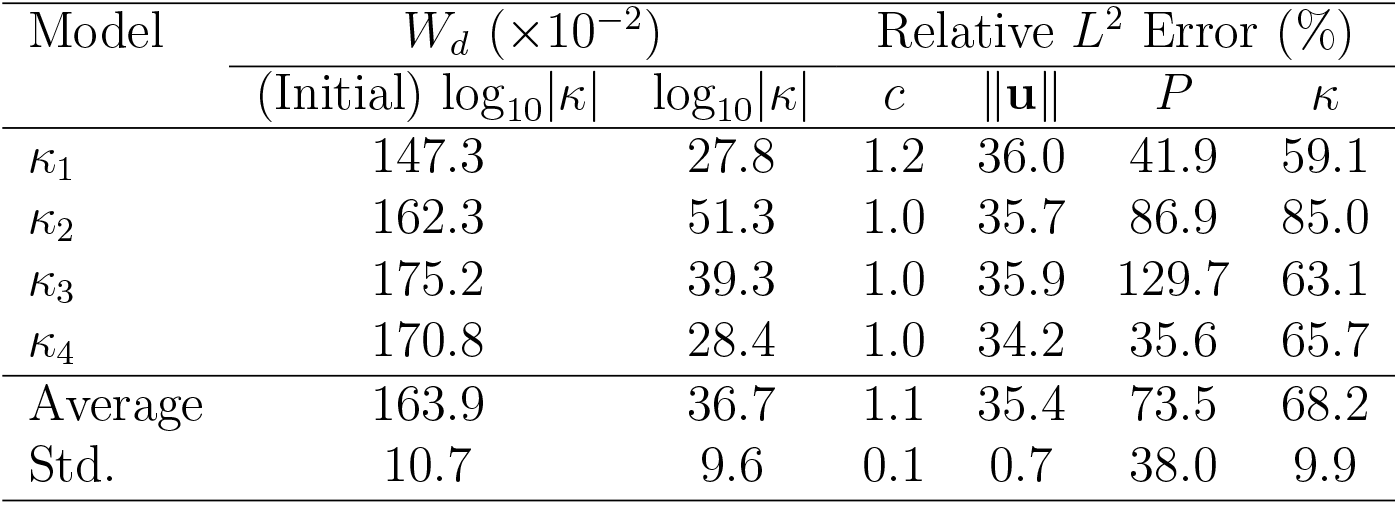
Permeability-based sensitivity analysis. The model shows strong and consistent performance in the velocity components, with relative errors in the velocity magnitude ∥**u**∥ varying by less than 2% across all cases. For pressure and permeability, the model can achieve low errors, but the uncertainty is substantially higher due to the ill-posed nature of the inverse problem under Darcy’s law. As a result, some initializations lead to accurate predictions while others do not. Nonetheless, the model consistently captures the overall distribution of the permeability and substantially reduces the Wasserstein distance (*W*_*d*_) between the log-scaled initial guess *κ* and the final prediction *κ*.

#### Appendix C.2.1. Velocity Inference

##### Synthetic Data

Figure C.14(A) shows the distribution of the predicted velocity components for models trained using Darcy’s law under different initial permeability guesses *κ*_1_–*κ*_4_. The model captures a wide dynamic range, particularly for the *w* component, which spans from 10^−6^ *µ*m*/*s to 10^3^ *µ*m*/*s. Additionally, the predictions consistently recover a bimodal distribution across all components, indicating robust structural consistency despite variations in permeability initialization. For comparison, we also show the velocity distribution from a baseline model that does not encode Darcy’s law in this architecture and predicts the velocity components directly. In this case, the advection-diffusion equation and mass conservation are insufficient to constrain the four unknowns *c, u, v*, and *w*, leading the model to collapse to a unimodal, average-like solution.

Table C.7 reports the relative *L*^2^ errors for each velocity component across permeability initializations. The model exhibits consistent performance, with an average relative *L*^2^ error of 35.4%. The variation across models is small, with standard deviations below 2% for all components.

Finally, to visualize the uncertainty distribution from different permeability initializations, Figure 7(A) shows the mean velocity magnitude 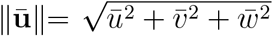 computed from the mean velocity components 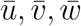 calculated from the ensemble average; see Equation C.1. Similarly, the figure also shows the relative uncertainty, computed as 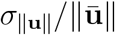, where *σ*_∥**u**∥_ denotes the standard deviation of the velocity magnitude across the ensemble. The bottom row presents the point-wise relative error with respect to the ground-truth velocity magnitude ∥**û**∥, computed as 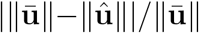. We observe that regions of high uncertainty often coincide with regions of high error, particularly near sharp permeability interfaces. Additionally, both error and uncertainty are elevated in low-velocity regions, where reconstruction is more sensitive to the initial guess. To quantify this observation, Figure C.14(C) shows the joint distributions of relative error vs. average velocity magnitude, relative error vs. relative uncertainty, and relative uncertainty vs. velocity magnitude. Notably, relative errors are concentrated in low-velocity regions, and the predicted uncertainty strongly correlates with true error. Relative errors can fall below 10% in regions of higher velocity, and overall, the relative error remains below 50% across most of the domain, indicating stable velocity reconstruction despite variations in permeability initialization.

##### Real Data

Figure C.14(B) shows the distribution of the predicted velocity components for Mouse 1 using different permeability initial guesses. As in the synthetic case, the predicted velocities exhibit a bimodal structure, with peaks at low and high velocities. While the high-velocity peak is consistent across models, there is some variability in the location and height of the low-velocity peak, typically centred around 10^−2^ *µ*m*/*s. In contrast, the baseline model, which does not enforce Darcy’s law, predicts a unimodal distribution, again indicating a collapse toward an average behaviour due to the lack of physical constraints. Overall, the ensemble captures a wide range of velocities spanning over four orders of magnitude, demonstrating the model’s flexibility and ability to resolve heterogeneous flow regimes even in the absence of ground-truth velocity data.

Figure 7(B) maps the mean velocity magnitude and its associated relative uncertainty for Mouse 1. Although we cannot compute point-wise errors in the real data due to the lack of a reference solution, the uncertainty map provides insight into prediction reliability. From the synthetic case, we observed that high uncertainty typically aligns with regions of low velocity, which are more sensitive to permeability initialization. A similar pattern emerges here: uncertainty is elevated in low-flow regions but remains below 100% across most of the domain. In contrast, uncertainty is low in regions with high velocity, suggesting that the model’s predictions are more reliable in those areas, even under varying initialization. Even though the relative uncertainty is large (uncertainty between 50 and 100% for over half of the brain), we consider this level of uncertainty acceptable, given that velocities in the deep brain were previously so uncertain.

#### Appendix C.2.2. Permeability and Pressure estimates

##### Synthetic Data

Table C.7 summarizes the variability in pressure and permeability reconstructions across different initializations. In contrast to the velocity components, which exhibit stable performance regardless of the starting point, both pressure and permeability are notably more sensitive to initialization. This behaviour reflects the ill-posed nature of the inverse problem under Darcy’s law, where multiple fields can satisfy the observed velocity data. As a result, some initializations yield relatively accurate estimates (e.g., *κ*_3_ with 35.6% error in pressure and 65.7% in permeability), while others lead to substantially higher errors. The elevated standard deviations in pressure (43.4%) and permeability errors (14.0%) underscore this inconsistency.

**Table D.8:** MR-AIV Architectures. MR-AIV Sub-networks number of layers and neurons per layer.

| Network | Layer | Neurons |
| --- | --- | --- |
| $NN_k$ | 5 | 150 |
| $NN_p$ | 5 | 150 |
| $NN_c$ | 5 | 200 |
| $NN_\sigma$ | 5 | 66 |

Despite this variability, all inferred permeability fields consistently reduce the mismatch between the initial guess (in log scale) and the final prediction, as measured by the Wasserstein distance. This distance provides a meaningful assessment of structural similarity between the predicted and true fields beyond point-wise errors. Spatial maps of the average predictions, relative uncertainties, and errors are presented in Fig. C.15 for permeability and pressure.

##### Real Data

The in-vivo results exhibit similar qualitative behaviour. As shown in Fig. C.15, uncertainty is primarily concentrated in regions with strong anatomical transitions and steep gradients. Notably, Fig. C.14 (D) shows that the predicted pressure and permeability remain surprisingly consistent across different initial guesses. This resemblance should not be misinterpreted as evidence of uniqueness. Rather, it may reflect the inherent complexity and spatial richness of the underlying field, which can effectively constrain the space of admissible solutions. Nevertheless, due to the ill-posed nature of the inverse problem, these reconstructions should be interpreted as plausible and physically consistent estimates, not as definitive ground-truth representations.

## Appendix D. Implementation Details

### Appendix D.1. Architectures

As described in Figure 6, the proposed method is composed of 4 Networks, and their architecture is described in Table D.8.

All models use the hyperbolic tangent as an activation function, so their inputs are normalized between (−1, 1) for stability [61].

### Appendix D.2. Training

For our experiments, development, training, validation, and visualization, we used JAX 0.3.2.9 as a machine learning framework on a single Nvidia A100-80GB GPU.

All models were trained using four AdamW [64] optimizers (i.e., one per network) with a weight decay of 10^−5^ for regularization. The initial and final learning rates are 2 × 10^−3^ and 5 × 10^−5^, respectively, with a decay rate of 0.99.

**Figure C.15:**
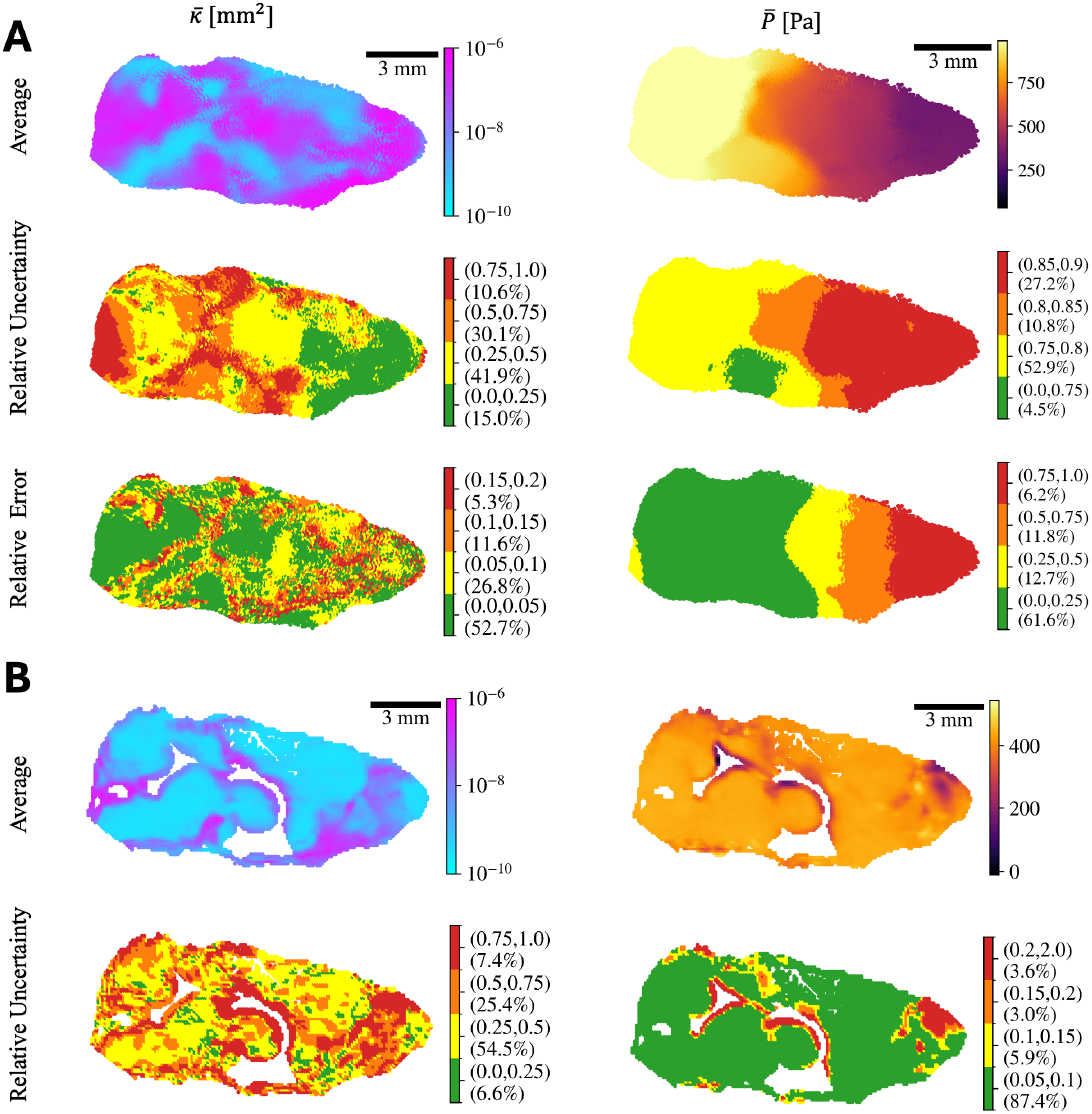
Uncertainty quantification of the inferred permeability and pressure fields. **(A)** (Top row) Average permeability 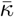 and pressure 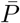 computed from four models with different initial permeability guesses, for the “realistic” synthetic data. (Middle row) Relative uncertainty across the four models. Colors are marked with the percentage of total brain volume to which each applies. (Bottom row) Relative error. Uncertainty is high around transition regions, which typically correspond to interfaces between low and high permeability zones. The permeability relative error reflects similar spatial trends and remains below 20% and 25% for permeability and pressure in most of the domain. **(B)** (Top row) Average permeability and pressure computed from four models with different initial permeability guesses, for Mouse 1. (Bottom row) Permeability relative uncertainty is highest near structural transitions and regions of elevated spatial variation, as with synthetic data. Pressure relative uncertainty is high near sharp gradients, as with the synthetic data, but is low in most of the domain.

**Table D.9:** Initialization stages: Global Weights (Step 0 and Step 1). In the initialization stages, the model follows a “data-driven” approach to initialize the concentration and permeability networks. For both initialization stages, we use an *L*_2_ norm loss for each loss sub-term *q* = 2.

| Stage | Loss Term | Sub-term | Global Weight ($m_\alpha$) |
| --- | --- | --- | --- |
| 0 | Data Low | $K$ | 100 |
| 0 | Data High | $K$ | 1 |
| 1 | Data | $c$ | 100 |
| 1 | Data | $\sigma_c$ | 100 |
| 1 | Initialization | $u, v, w$ | 100 |
| 1 | Equations | $AD$ (see equation A.9) | $10^{-16}$ |
| 1 | Equations | $CM$ (see equation A.10) | $10^{-16}$ |

**Table D.10:** Training stages: Global Weights (Step 2 and Step 3). Step 2 updates only the pressure network parameters (***θ***_*P*_). Step 3 updates parameters for both pressure (***θ***_*P*_) and permeability (***θ***_*K*_). Step 4 introduces concentration updates, modifying parameters for pressure (***θ***_*P*_), permeability (***θ***_*K*_), and concentration (***θ***_*c*_). Finally, Step 5 continues updating all parameters (***θ***_*P*_, ***θ***_*K*_, ***θ***_*c*_) but utilizes the *L*^1^ norm (*q* = 1).

| Stage | Loss Term | Sub-term | Global Weight ($m_\alpha$) |
| --- | --- | --- | --- |
| 2 | Equations | $AD$ (see equation A.9) | 10 |
| 2 | Equations | $CM$ (see equation A.10) | $10^{-2}$ |
| 3 | Equations | $AD$ (see equation A.9) | 10 |
| 3 | Equations | $CM$ (see equation A.10) | $10^{-2}$ |
| 4 | Data | $c$ | 100 |
| 4 | Equations | $AD$ (see equation A.9) | 10 |
| 4 | Equations | $CM$ (see equation A.10) | $10^{-2}$ |
| 5 | Data | $c$ | 10 |
| 5 | Equations | $AD$ (see equation A.9) | 10 |
| 5 | Equations | $CM$ (see equation A.10) | $10^{-2}$ |

For the RBA-R hyperparameters, we follow [37] and use *γ* = 0.999, *η* = 0.1, *ν* = 2.0, and *c* = 0.5 (see equation A.20). All models were trained using a batch size of *bs* = 10000.

#### Appendix D.2.1. Sequential Training

In all cases, the sequential strategy was imposed by varying the global weights *m*_*α*_ during the initialization and training stages as described in Tables D.10 and D.9, respectively.

## References

[1] L. Xie, H. Kang, Q. Xu, M. J. Chen, Y. Liao, M. Thiyagarajan, J. O’Donnell, D. J. Christensen, C. Nicholson, J. J. Iliff, T. Takano, R. Deane, M. Nedergaard, Sleep drives metabolite clearance from the adult brain, Science 342 (6156) (2013) 373–377.

[2] M. K. Rasmussen, H. Mestre, M. Nedergaard, The glymphatic pathway in neuro-logical disorders, The Lancet Neurology 17 (2018) 1016–1024, doi: 10.1016/S1474-4422(18)30318-1. doi:10.1016/S1474-4422(18)30318-1.

[3] B. A. Plog, M. Nedergaard, The Glymphatic System in Central Nervous System Health and Disease: Past, Present, and Future, Annu. Rev. Pathol. Mech. Dis. 13 (1) (2018) annurev–pathol–051217–111018–19.

[4] H. Mestre, T. Du, A. M. Sweeney, G. Liu, A. J. Samson, W. Peng, K. N. Mortensen, F. F. Stæger, P. A. R. Bork, L. Bashford, E. R. Toro, J. Tithof, D. H. Kelley, J. H. Thomas, P. G. Hjorth, E. A. Martens, R. I. Mehta, O. Solis, P. Blinder, D. Kleinfeld, H. Hirase, Y. Mori, M. Nedergaard, Cerebrospinal fluid influx drives acute ischemic tissue swelling, Science 9 (2020) eaax7171–24.

[5] D. H. Kelley, J. H. Thomas, Cerebrospinal fluid flow, Annual Review of Fluid Mechanics 55 (1) (2023) 237–264.

[6] H. Mestre, J. Tithof, T. Du, W. Song, W. Peng, A. M. Sweeney, G. Olveda, J. H. Thomas, M. Nedergaard, D. H. Kelley, Flow of cerebrospinal fluid is driven by arterial pulsations and is reduced in hypertension, Nat. Commun. 9 (1) (2018) 4878.

[7] F. Min Rivas, J. Liu, B. C. Martell, T. Du, H. Mestre, M. Nedergaard, J. Tithof, J. H. Thomas, D. H. Kelley, Surface periarterial spaces of the mouse brain are open, not porous, J. R. Soc. Interface 17 (172) (2020) 20200593.

[8] A. Raghunandan, A. Ladron-de Guevara, J. Tithof, H. Mestre, T. Du, M. Nedergaard, J. H. Thomas, D. H. Kelley, Bulk flow of cerebrospinal fluid observed in periarterial spaces is not an artifact of injection, Elife 10 (2021) e65958.

[9] K. A. Boster, S. Cai, A. Ladrón-de Guevara, J. Sun, X. Zheng, T. Du, J. H. Thomas, M. Nedergaard, G. E. Karniadakis, D. H. Kelley, Artificial intelligence velocimetry reveals in vivo flow rates, pressure gradients, and shear stresses in murine perivascular flows, Proceedings of the National Academy of Sciences 120 (14) (2023) e2217744120.

[10] J. D. Toscano, C. Wu, A. Ladrón-de Guevara, T. Du, M. Nedergaard, D. H. Kelley, G. E. Karniadakis, K. A. Boster, Inferring in vivo murine cerebrospinal fluid flow using artificial intelligence velocimetry with moving boundaries and uncertainty quantification, Interface Focus 14 (6) (2024) 20240030.

[11] B. A. Plog, H. Mestre, G. E. Olveda, A. M. Sweeney, H. M. Kenney, A. Cove, K. Y. Dholakia, J. Tithof, T. D. Nevins, I. Lundgaard, T. Du, D. H. Kelley, M. Nedergaard, Transcranial optical imaging reveals a pathway for optimizing the delivery of immunotherapeutics to the brain, JCI Insight 3 (20) (2018) 1188–16.

[12] A. S. Munk, W. Wang, N. B. Bèchet, A. M. Eltanahy, A. X. Cheng, B. Sigurdsson, A. Benraiss, M. A. Mäe, B. T. Kress, D. H. Kelley, C. Betsholtz, K. Møllgård, A. Meissner, M. Nedergaard, I. Lundgaard, PDGF-B Is Required for Development of the Glymphatic System, Cell Rep 26 (11) (2019) 2955–2969.e3.

[13] L. M. Valnes, S. K. Mitusch, G. Ringstad, P. K. Eide, S. W. Funke, K.-A. Mardal, Apparent diffusion coefficient estimates based on 24 hours tracer movement support glymphatic transport in human cerebral cortex, Scientific Reports 10 (2020) 9176. doi:10.1038/s41598-020-66042-5. URL https://doi.org/10.1038/s41598-020-66042-5

[14] L. A. Ray, M. Pike, M. Simon, J. J. Iliff, J. J. Heys, Quantitative analysis of macroscopic solute transport in the murine brain, Fluids and Barriers of the CNS 18 (2021) 55. doi:10.1186/s12987-021-00290-z. URL https://doi.org/10.1186/s12987-021-00290-z

[15] B. Zapf, J. Haubner, M. Kuchta, G. Ringstad, P. K. Eide, K.-A. Mardal, Investigating molecular transport in the human brain from mri with physics-informed neural networks, Scientific Reports 12 (1) (2022) 15475.

[16] J. Oldenburg, J. Renkewitz, M. Stiehm, K.-P. Schmitz, Contributions towards data driven deep learning methods to predict steady state fluid flow in mechanical heart valves, Current Directions in Biomedical Engineering 7 (2) (2021) 625–628 [cited 2025-02-17]. doi:10.1515/cdbme-2021-2159. URL https://doi.org/10.1515/cdbme-2021-2159

[17] X. Hou, P. Guo, P. Wang, P. Liu, D. D. Lin, H. Fan, Y. Li, Z. Wei, Z. Lin, D. Jiang, et al., Deep-learning-enabled brain hemodynamic mapping using restingstate fmri, npj Digital Medicine 6 (1) (2023) 116.

[18] D. R. Rutkowski, A. Roldán-Alzate, K. M. Johnson, Enhancement of cerebrovascular 4d flow mri velocity fields using machine learning and computational fluid dynamics simulation data, Scientific reports 11 (1) (2021) 10240.

[19] S. Talebi, S. Gai, A. Sossin, V. Zhu, E. Tong, M. R. Mofrad, Deep learning for perfusion cerebral blood flow (cbf) and volume (cbv) predictions and diagnostics, Annals of Biomedical Engineering 52 (6) (2024) 1568–1575.

[20] V. Ratner, Y. Gao, H. Lee, R. Elkin, M. Nedergaard, H. Benveniste, A. Tannenbaum, Cerebrospinal and interstitial fluid transport via the glymphatic pathway modeled by optimal mass transport, NeuroImage 152 (2017) 530–537.

[21] A. F. Frangi, J. A. Schnabel, C. Davatzikos, C. Alberola-López, G. Fichtinger (Eds.), GlymphVIS: Visualizing Glymphatic Transport Pathways Using Regularized Optimal Transport, Springer International Publishing, Cham, 2018.

[22] S. Koundal, R. Elkin, S. Nadeem, Y. Xue, S. Constantinou, S. Sanggaard, X. Liu, B. Monte, F. Xu, W. Nostrand, M. Nedergaard, H. Lee, J. Wardlaw, H. Benveniste, A. Tannenbaum, Optimal Mass Transport with Lagrangian Workflow Reveals Advective and Diffusion Driven Solute Transport in the Glymphatic System, Sci. Rep. (2020) 1–18.

[23] X. Chen, X. Liu, S. Koundal, R. Elkin, X. Zhu, B. Monte, F. Xu, F. Dai, M. Pedram, H. Lee, J. Kipnis, A. Tannenbaum, W. E. V. Nostrand, H. Benveniste, Cerebral amyloid angiopathy is associated with glymphatic transport reduction and time-delayed solute drainage along the neck arteries, Nature Aging 2 (2022) 214–223. doi:10.1038/s43587-022-00181-4. URL https://doi.org/10.1038/s43587-022-00181-4

[24] S. Koundal, X. Chen, Z. Gursky, H. Lee, K. Xu, F. Liang, Z. Xie, F. Xu, H.-M. Lin, W. E. V. Nostrand, X. Gu, R. Elkin, A. Tannenbaum, H. Benveniste, Divergent brain solute clearance in rat models of cerebral amyloid angiopathy and alzheimer’s disease, iScience 27, doi: 10.1016/j.isci.2024.111463 (12 2024). doi:10.1016/j.isci.2024.111463. URL https://doi.org/10.1016/j.isci.2024.111463

[25] V. Vinje, B. Zapf, G. Ringstad, P. K. Eide, M. E. Rognes, K.-A. Mardal, Human brain solute transport quantified by glymphatic mri-informed biophysics during sleep and sleep deprivation, Fluids and Barriers of the CNS 20 (2023) 62. doi: 10.1186/s12987-023-00459-8. URL https://doi.org/10.1186/s12987-023-00459-8

[26] T. Bohr, P. G. Hjorth, S. C. Holst, S. Hrabětová, V. Kiviniemi, T. Lilius, I. Lundgaard, K.-A. Mardal, E. A. Martens, Y. Mori, U. V. Nägerl, C. Nicholson, A. Tannenbaum, J. H. Thomas, J. Tithof, H. Benveniste, J. J. Iliff, D. H. Kelley, M. Nedergaard, The glymphatic system: Current understanding and modeling, iScience 25 (9) (2022) 104987.

[27] L. Ray, J. J. Iliff, J. J. Heys, Analysis of convective and diffusive transport in the brain interstitium, Fluids Barriers CNS 16 (1) (2019) 6.

[28] F. Min Rivas, J. Liu, B. C. Martell, T. Du, H. Mestre, M. Nedergaard, J. Tithof, J. H. Thomas, D. H. Kelley, Surface periarterial spaces of the mouse brain are open, not porous, Journal of The Royal Society Interface 17 (172) (2020) 20200593. arXiv:https://royalsocietypublishing.org/doi/pdf/10.1098/rsif.2020.0593, doi:10.1098/rsif.2020.0593. URL https://royalsocietypublishing.org/doi/abs/10.1098/rsif.2020.0593

[29] K. E. Holter, B. Kehlet, A. Devor, T. J. Sejnowski, A. M. Dale, S. W. Omholt, O. P. Ottersen, E. A. Nagelhus, K.-A. Mardal, K. H. Pettersen, Interstitial solute transport in 3D reconstructed neuropil occurs by diffusion rather than bulk flow, Proc. Nat. Acad. Sci. 114 (37) (2017) 9894–9899.

[30] M. K. Sharp, R. O. Carare, B. A. Martin, Dispersion in porous media in oscillatory flow between flat plates: applications to intrathecal, periarterial and paraarterial solute transport in the central nervous system, Fluids and Barriers of the CNS 16 (2019) 1–17.

[31] F. Romanò, V. Suresh, P. A. Galie, J. B. Grotberg, Peristaltic flow in the glymphatic system, Sci. Rep. (2020) 1–17.

[32] R. T. Kedarasetti, P. J. Drew, F. Costanzo, Arterial vasodilation drives convective fluid flow in the brain: a poroelastic model, Fluids and Barriers of the CNS 19 (1) (2022) 34.

[33] M. M. Meerschaert, C. Tadjeran, Finite difference approximations for fractional advection–dispersion flow equations, Journal of Computational and Applied Mathematics 172 (1) (2004) 65–77.

[34] M. Raissi, P. Perdikaris, G. E. Karniadakis, Physics-informed neural networks: A deep learning framework for solving forward and inverse problems involving nonlinear partial differential equations, Journal of Computational Physics 378 (2019) 686–707.

[35] M. Raissi, A. Yazdani, G. E. Karniadakis, Hidden Fluid Mechanics: A Navier-Stokes Informed Deep Learning Framework for Assimilating Flow Visualization Data, arXiv preprint 1808.04327 (2018).

[36] S. Cai, H. Li, F. Zheng, F. Kong, M. Dao, G. E. Karniadakis, S. Suresh, Artificial intelligence velocimetry and microaneurysm-on-a-chip for three-dimensional analysis of blood flow in physiology and disease, Proceedings of the National Academy of Sciences 118 (13) (2021).

[37] J. D. Toscano, T. Käufer, Z. Wang, M. Maxey, C. Cierpka, G. E. Karniadakis, Aivt: Inference of turbulent thermal convection from measured 3d velocity data by physics-informed kolmogorov-arnold networks, Science Advances 11 (19) (2025) eads5236.

[38] S. Kida, R. O. Weller, E.-T. Zhang, M. J. Phillips, F. Iannotti, Anatomical pathways for lymphatic drainage of the brain and their pathological significance, Neuropathology and Applied Neurobiology 21 (3) (1995) 181–184. arXiv:https://onlinelibrary.wiley.com/doi/pdf/10.1111/j.1365-2990.1995.tb01048.x, doi:10.1111/j.1365-2990.1995.tb01048.x. URL https://onlinelibrary.wiley.com/doi/abs/10.1111/j.1365-2990.1995.tb01048.x

[39] H. Mansour, R. Azrak, J. J. Cook, K. J. Hornburg, Y. Qi, Y. Tian, R. W. Williams, F.-C. Yeh, L. E. White, G. A. Johnson, The duke mouse brain atlas: Mri and light sheet microscopy stereotaxic atlas of the mouse brain, Science Advances 11 (18) (2025) eadq8089.

[40] K. A. S. Boster, J. Tithof, D. D. Cook, J. H. Thomas, D. H. Kelley, Sensitivity analysis on a network model of glymphatic flow, Journal of The Royal Society Interface 19 (191) (2022) 20220257. arXiv: https://royalsocietypublishing.org/doi/pdf/10.1098/rsif.2022.0257, doi:10.1098/rsif.2022.0257. URL https://royalsocietypublishing.org/doi/abs/10.1098/rsif.2022.0257

[41] B. Lakshminarayanan, A. Pritzel, C. Blundell, Simple and scalable predictive uncertainty estimation using deep ensembles, Advances in Neural Information Processing Systems 30 (2017).

[42] S. J. Anagnostopoulos, J. D. Toscano, N. Stergiopulos, G. E. Karniadakis, Learning in PINNs: Phase transition, total diffusion, and generalization, arXiv preprint 2403.18494 (2024).

[43] S. J. Anagnostopoulos, J. D. Toscano, N. Stergiopulos, G. E. Karniadakis, Residual-based attention in physics-informed neural networks, Computer Methods in Applied Mechanics and Engineering 421 (2024) 116805.

[44] J. D. Toscano, L.-L. Wang, G. E. Karniadakis, Kkans: Kurkova-kolmogorov-arnold networks and their learning dynamics, Neural Networks (2025) 107831.

[45] K. Shukla, J. D. Toscano, Z. Wang, Z. Zou, G. E. Karniadakis, A comprehensive and FAIR comparison between MLP and KAN representations for differential equations and operator networks, Computer Methods in Applied Mechanics and Engineering 431 (2024) 117290.

[46] C. Wu, J. D. Toscano, K. Shukla, Y. Chen, A. Shahmohammadi, E. Raymond, T. Toupy, N. Nazemifard, C. Papageorgiou, G. E. Karniadakis, Fmenets: Flow, material, and energy networks for non-ideal plug flow reactor design, arXiv preprint 2505.20300 (2025).

[47] J. J. Iliff, M. Wang, Y. Liao, B. A. Plog, W. Peng, G. A. Gundersen, H. Benveniste, G. E. Vates, R. Deane, S. A. Goldman, E. A. Nagelhus, M. Nedergaard, A Paravascular Pathway Facilitates CSF Flow Through the Brain Parenchyma and the Clearance of Interstitial Solutes, Including Amyloid, Science Translational Medicine 4 (147) (2012) 147ra111–147ra111.

[48] J. V. George, K. J. Hornburg, A. Merrill, E. Marvin, K. Conrad, K. Welle, R. Gelein, D. Chalupa, U. Graham, G. Oberdörster, et al., Brain iron accumulation in neurodegenerative disorders: Does air pollution play a role?, Particle and Fibre Toxicology 22 (1) (2025) 9.

[49] R. D. Penn, A. Linninger, The physics of hydrocephalus, Pediatric Neurosurgery 45 (3) (2009) 161–174.

[50] Y. Guo, K. Quirk, D. H. Kelley, J. H. Thomas, Advection and diffusion in perivascular and extracellular spaces in the brain, Journal of The Royal Society Interface 22 (226) (2025) 20250010.

[51] B. Bedussi, M. Almasian, J. de Vos, E. VanBavel, E. N. Bakker, Paravascular spaces at the brain surface: Low resistance pathways for cerebrospinal fluid flow, Journal of cerebral blood flow and metabolism : official journal of the International Society of Cerebral Blood Flow and Metabolism 38 (2018) 719–726. doi:10.1177/0271678X17737984. URL https://pubmed.ncbi.nlm.nih.gov/nih.gov/pmc/articles/PMC5888857/

[52] A. Raghunandan, A. L. de Guevara, J. Tithof, H. Mestre, T. Du, M. Nedergaard, J. H. Thomas, D. H. Kelley, Bulk flow of cerebrospinal fluid observed in periarterial spaces is not an artifact of injection, eLife 10 (2021) e65958. doi:10.7554/eLife.65958. URL https://doi.org/10.7554/eLife.65958

[53] P. J. Basser, Interstitial pressure, volume, and flow during infusion into brain tissue, Microvasc. Res. 44 (2) (1992) 143–165.

[54] G. A. Johnson, Y. Tian, D. G. Ashbrook, G. P. Cofer, J. J. Cook, J. C. Gee, A. Hall, K. Hornburg, C. C. Kaczorowski, Y. Qi, et al., Merged magnetic resonance and light sheet microscopy of the whole mouse brain, Proceedings of the National Academy of Sciences 120 (17) (2023) e2218617120.

[55] X. Han, S. Maharjan, J. Chen, Y. Zhao, Y. Qi, L. E. White, G. A. Johnson, N. Wang, High-resolution diffusion magnetic resonance imaging and spatialtranscriptomic in developing mouse brain, Neuroimage 297 (2024) 120734.

[56] E. H. Stanton, N. D. Åke Persson, R. S. Gomolka, T. Lilius, B. Sigurdsson, H. Lee, A. L. R. Xavier, H. Benveniste, M. Nedergaard, Y. Mori, Mapping of csf transport using high spatiotemporal resolution dynamic contrast-enhanced mri in mice: Effect of anesthesia, Magnetic Resonance in Medicine 85 (2021) 3326–3342. doi:10.1002/mrm.28645. URL https://doi.org/10.1002/mrm.28645

[57] R. S. Gomolka, L. M. Hablitz, H. Mestre, M. Giannetto, T. Du, N. L. Hauglund, L. Xie, W. Peng, P. M. Martinez, M. Nedergaard, Y. Mori, Loss of aquaporin-4 results in glymphatic system dysfunction via brain-wide interstitial fluid stagnation, eLife 12 (2023) e82232. doi:10.7554/eLife.82232. URL >https://doi.org/10.7554/eLife.82232

[58] T. D. Nevins, D. H. Kelley, Front tracking for quantifying advection-reactiondiffusion, Chaos 27 (4) (2017) 043105–10.

[59] T. D. Nevins, D. H. Kelley, Front tracking velocimetry in advection-reaction-diffusion systems, Chaos 28 (4) (2018) 043122–11.

[60] G. Ringstad, L. M. Valnes, A. M. Dale, A. H. Pripp, S.-A. S. Vatnehol, K. E. Emblem, K.-A. Mardal, P. K. Eide, Brain-wide glymphatic enhancement and clearance in humans assessed with mri, JCI Insight 3 (13) (7 2018). doi: 10.1172/jci.insight.121537. URL https://doi.org/10.1172/jci.insight.121537

[61] J. D. Toscano, V. Oommen, A. J. Varghese, Z. Zou, N. Ahmadi Daryakenari, C. Wu, G. E. Karniadakis, From pinns to pikans: Recent advances in physics-informed machine learning, Machine Learning for Computational Science and Engineering 1 (1) (2025) 1–43.

[62] T. Salimans, D. P. Kingma, Weight normalization: A simple reparameterization to accelerate training of deep neural networks, Advances in Neural Information Processing Systems 29 (2016).

[63] S. Wang, B. Li, Y. Chen, P. Perdikaris, PirateNets: Physics-informed Deep Learning with Residual Adaptive Networks, arXiv preprint 2402.00326 (2024).

[64] I. Loshchilov, F. Hutter, Decoupled weight decay regularization, arXiv preprint 1711.05101 (2017).

[65] L. D. McClenny, U. M. Braga-Neto, Self-adaptive physics-informed neural networks, Journal of Computational Physics 474 (2023) 111722.

[66] C. Wu, M. Zhu, Q. Tan, Y. Kartha, L. Lu, A comprehensive study of non-adaptive and residual-based adaptive sampling for physics-informed neural networks, Computer Methods in Applied Mechanics and Engineering 403 (2023) 115671.

[67] A. F. Psaros, X. Meng, Z. Zou, L. Guo, G. E. Karniadakis, Uncertainty quantification in scientific machine learning: Methods, metrics, and comparisons, Journal of Computational Physics 477 (2023) 111902.

[68] Z. Zou, X. Meng, G. E. Karniadakis, Uncertainty quantification for noisy inputsoutputs in physics-informed neural networks and neural operators, arXiv preprint 2311.11262 (2023).

[69] Z. Zou, X. Meng, A. F. Psaros, G. E. Karniadakis, NeuralUQ: A comprehensive library for uncertainty quantification in neural differential equations and operators, SIAM Review 66 (1) (2024) 161–190.

